# LipidMS 3.0: an R-package and a web-based tool for LC-MS/MS data processing and lipid annotation

**DOI:** 10.1101/2022.02.25.476005

**Authors:** María Isabel Alcoriza-Balaguer, Juan Carlos García-Cañaveras, Francisco Javier Ripoll-Esteve, Marta Roca, Agustín Lahoz

## Abstract

**Summary:** LipidMS was initially envisioned to use fragmentation rules and data-independent acquisition (DIA) for lipid annotation. However, data-dependent acquisition (DDA) remains the most widespread acquisition mode for untargeted LC-MS/MS-based lipidomics. Here we present LipidMS 3.0, an R package that not only adds DDA and new lipid classes to its pipeline, but also the required functionalities to cover the whole data analysis workflow from pre-processing (i.e., peak-peaking, alignment and grouping) to lipid annotation. We applied the new workflow in the data analysis of a commercial human serum pool spiked with 68 lipid standards acquired in the full scan, DDA and DIA modes. When focusing on the detected lipid standard features and total identified lipids, LipidMS 3.0 data pre-processing performance is similar to XCMS, whereas it complements the annotations provided by MS-DIAL, one of the most widely used tools in lipidomics. To extend and facilitate LipidMS 3.0 usage among less experienced R-programming users, the workflow was also implemented as a web-based application.

**Availability and Implementation:** The LipidMS R-package is freely available at https://CRAN.R-project.org/package=LipidMS and as a website at http://www.lipidms.com.

## 1 Introduction

Lipids are a heterogeneous group of non polar molecules that play important biological roles, such as structural components of cell membranes, energy storage or signaling molecules (Wenk, 2005). Recent advances made in mass spectrometry (MS)-based analyses of the lipidome, lipid annotation tools and *in silico*-generated spectra databases have enabled the characterized lipidome to continuously expand (Züllig *et al*., 2020). Liquid chromatography (LC) coupled with high-resolution mass spectrometry is the most prominent analytical platform for untargeted lipidomics (Roca *et al*., 2021). Currently, there are two main approaches to generate LC-MS/MS data: data-dependent acquisition (DDA), in which precursors from MS1 (full-scan) are selected and immediately fragmented (MS2) to generate a spectrum that can be directly queried against a spectra database (Koelmel, Kroeger, Gill, *et al*., 2017); data-independent acquisition (DIA), where no precursor from MS1 is isolated and all the ions are subsequently fragmented (Guo and Huan, 2020). A wide range of tools has been developed for lipid annotation based on both DDA and DIA (Koelmel, Kroeger, Ulmer, *et al*., 2017; Alcoriza-Balaguer *et al*., 2019; Tsugawa *et al*., 2015; Hartler *et al*., 2017). Of them, the LipidMS R package (Alcoriza-Balaguer *et al*., 2019) was envisioned to use fragmentation rules and DIA to annotate lipids with different structural elucidation levels. However, it required external software to process large datasets and missed the option of using DDA. Here we present LipidMS 3.0, a new R package release and web-based application (www.lipidms.com), which enables the use of DDA, adds new lipid classes to its repertoire, and includes new functionalities to cover the whole data processing workflow from pre-processing (i.e., peak-peaking, alignment and grouping) to lipid annotation (**Figure 1**).

**Figure 1.**
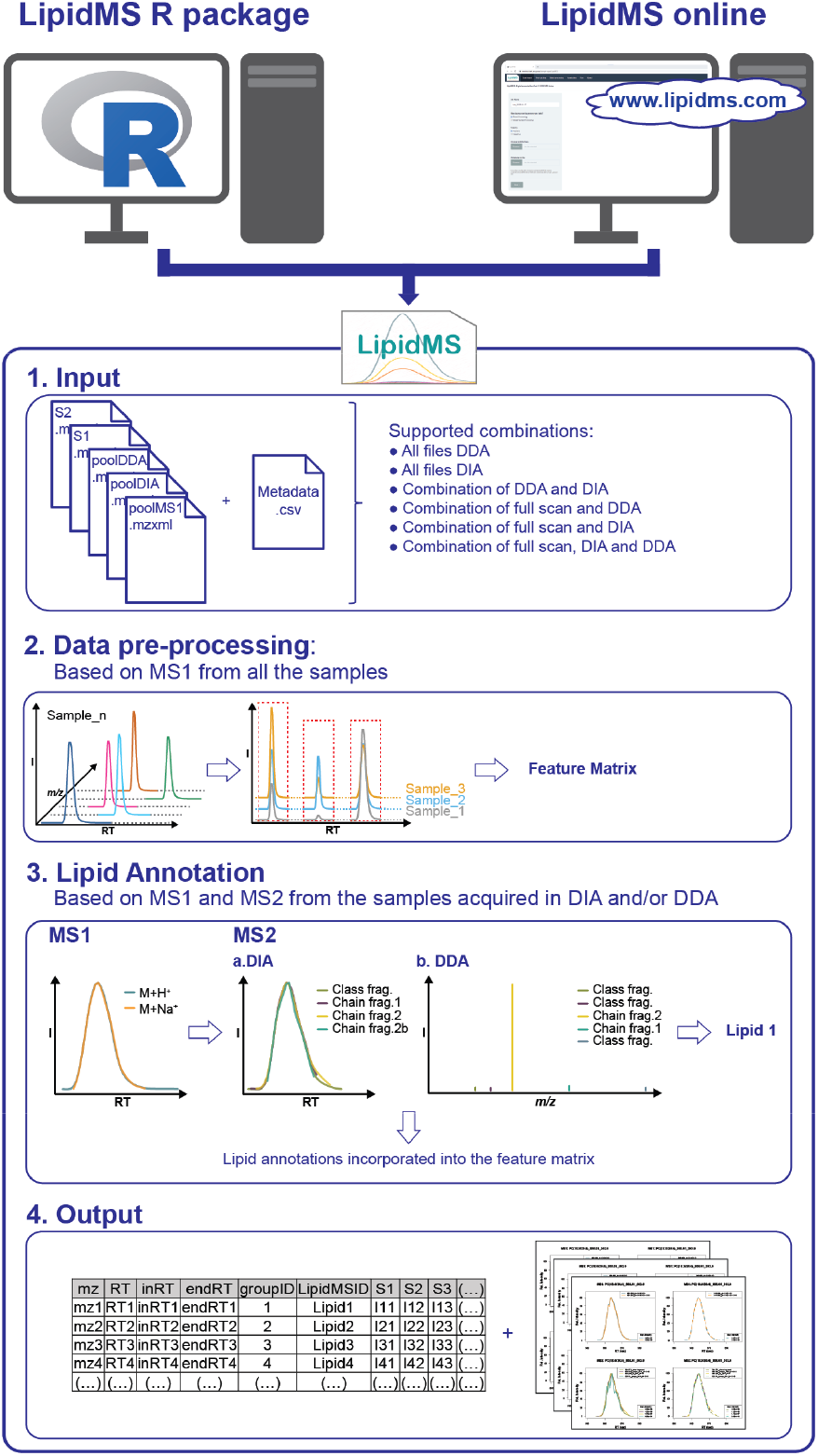
LipidMS 3.0 workflow. LipidMS is an open-access platform-independent software for lipid annotation from LC-MS/MS-based lipidomics that can be executed locally with an R environment (https://CRAN.R-project.org/package=LipidMS) or online (www.lipidms.com). Briefly, the LipidMS workflow comprises the following steps: 1) raw data files in the mzXML format and a csv metadata file (sample, acquisition mode, which can be full scan, DDA or DIA, and sample type) are used as input; 2) data pre-processing, including peak-peaking, alignment, grouping and peak filling, is executed based on the MS1 level information from all the samples, and a feature matrix is generated; 3) lipids are annotated based on the established fragmentation rules for those samples acquired in DIA or DDA, and the identifications incorporated into the feature matrix generated in step 2; 4) two main outputs are returned: a data matrix containing the peak areas and lipid annotations, if obtained, for all the features and samples found in the dataset, and plots showing the fragments that support the proposed lipid identifications.

To test LipidMS performance, a dataset acquired in the full scan, DDA and DIA modes, obtained upon the LC-MS lipidomic analysis of a commercial human serum pool spiked with 68 lipid standards, was used. LipidMS v3.0 data pre-processing was compared to XCMS 3.16 (Smith *et al*., 2006), whereas lipid annotation was compared to MS-DIAL 4.80 (Tsugawa *et al*., 2015), the current state of the art for lipidomics analysis.

## 2 Features and implementation

### 2.1 Data pre-processing

LipidMS was initially designed to annotate lipids in a single sample files and, to this end, external software was required for batch data pre-processing. In its new version, LipidMS 3.0 includes a battery of data pre-processing functions to analyze untargeted LC-MS and LC-MS/MS datasets:

- ***Peak-picking***. The first step in LipidMS pre-processing consists in extracting a peak list for each sample of the dataset using the enviPick algorithm (https://CRAN.R-project.org/package=enviPick). This function also includes annotation for the ^13^C isotopes based on mass difference, relative intensity and correlation between peaks.
- ***Peak alignment***. Once peak lists have been extracted, the recurrent peaks from different samples are grouped based on mass-to-charge ratio (*m/z*) and retention time (RT) clustering (**Figures S1 and S2**). Then LOESS regression is applied to correct RT drifts among samples.
- ***Peak grouping***. After samples alignment, the peaks from the different samples that correspond to the same feature are grouped based on *m/z* and RT by using the same clustering algorithms as in peak alignment (**Figures S1 and S2**), but by employing more restrictive parameters for *m/z* and RT tolerance. The result is a feature matrix for the dataset.
- ***Filling peaks***. Once all feature peaks have been defined, peak areas are extracted again by a target approach that avoids missing peaks and improves area estimations.

Further details of these pre-processing functions can be found in the **Supplementary Information**.

### 2.3 Data-dependent acquisition

Despite the important advancements made for DIA-based annotation, DDA remains the most widespread data acquisition mode in LC-MS/MS (Guo and Huan, 2020). For this reason, LipidMS 3.0 now incorporates both data acquisition methods. For DDA, the same previously established fragmentation rules are applied (Alcoriza-Balaguer *et al*., 2019), but now fragments are searched directly in MS/MS spectra linked with a specific precursor from MS1. For each precursor ion, the MS/MS spectra used for annotation are the closest to precursor RT, which improves differentiation between isomeric lipid species.

### 2.3 New lipid classes

LipidMS initially covered 24 classes, for which fragmentation rules were optimized based on experimental and *in silico*-based spectra (Alcoriza-Balaguer *et al*., 2019). Ever since it was first released, new fragmentation rules for six lipid classes [acylceramides (AcylCer), ceramides phosphate (CerP), plasmanyl and plasmenyl phosphocholines (PCp, PCo) and phosphoethanolamines (PEp, PEo)] have been obtained based on the experimental spectra of available commercial lipids. Hence these new lipid classes have been added to the LipidMS 3.0 repertoire (**Supplementary Tables S1-S3** and **Supplementary Figures S3-S14**).

### 2.4 LipidMS workflow

LipidMS 3.0 uses raw data files in the mzXML format, which can be obtained employing msConvert from ProteoWizard (Chambers *et al*., 2012), and a csv metadata file, which contains sample names, acquisition mode (full scan, DDA or DIA) and sample type (blank, quality control, group 1, etc.) as input (**Figure 1**). The recommended workflow to follow is to acquire samples in full scan and a pooled sample in full scan, and in DDA or DIA, which is utilized to perform lipid annotation. LipidMS 3.0 supports the simultaneous data processing of any combination of full scan, DIA and DDA. Data pre-processing is performed with the algorithms and parameters optimized for common lipidomics approaches in which the elution of multiple isomers in a narrow RT window is common (Alcoriza-Balaguer *et al*., 2019). Lipid identification is based on fragmentation rules using DIA (based on co-elution) or DDA (clean spectra) data, and finally the potential identities are matched to the whole dataset. LipidMS provides two outputs: a table that summarizes the intensity and identity of all the detected lipids across samples; plots depicting information that supports the proposed lipid identities, as well as the achieved level of confidence for each identification (Alcoriza-Balaguer *et al*., 2019).

### 2.5 Implementation

LipidMS 3.0 is developed as an R package and is available via CRAN (https://CRAN.R-project.org/package=LipidMS). The source code and development version are also available at https://github.com/maialba3/LipidMS. While the R environment presents several advantages, such as highly flexible and customizable workflows that allow the management of large datasets, it requires R programming skills to make the most of its usage. To access a broader range of users, LipidMS 3.0 has also been implemented as a web-based application with a user-friendly GUI interface (**Supplementary Figure S15**), which is accessible at http://www.lipidms.com. Example data files, scripts and tutorials for the R package and the web application can be found at http://www.lipidms.com via the “Resources” tab.

### 2.6 Performance Evaluation

To evaluate LipidMS 3.0 performance, a commercial human serum pool spiked with 68 lipid standards was analyzed by LC-MS/MS using both the ESI + and ESI – ionization modes and full scan, DIA and DDA acquisition modes.

First, LipidMS 3.0 data pre-processing performance was compared to XCMS 3.16, one of the most widely used tools for LC-MS data processing (Smith *et al*., 2006). Despite the fact that XCMS found a larger total number of features than LipidMS 3.0 (33352 vs. 19382) (**Supplementary Table S4**), both software provided comparable numbers in terms of expected lipid standard features detected (129 features were expected from the 68 lipid standards, 122 and 114 were extracted by LipidMS 3.0 and XCMS 3.16, respectively) (**Supplementary Tables S4-S6**).

Then the LipidMS 3.0 complete workflow performance was compared to MS-DIAL 4.80 (Tsugawa *et al*., 2015). MS-DIAL extracted a much larger number of features than LipidMS 3.0 (75263 vs. 25574) (**Supplementary Table S4**), but in both software, a similar number of expected lipid standard features was found (**Supplementary Tables S4-S6**). However, LipidMS outperformed MS-DIAL in identified features (98/129 vs. 76/129) and identified lipids (60/68 vs. 56/68) (**Supplementary Tables S4-S6**). Then all the annotations provided by MS-DIAL and LipidMS 3.0 were manually curated and the results compared. MS-DIAL provided a larger number of correctly annotated lipids than LipidMS 3.0 (580 vs. 387 in ESI – and 588 vs. 445 in ESI +) (**Supplementary Tables S7-S8**), but with a much larger number of incorrect annotations (669 vs. 79 in ESI – and 897 vs. 50 in ESI+) (**Supplementary Tables S7-S8**). So while the LipidMS incorrect annotations represented 10-17 % of the proposed identities, for MS-DIAL it accounted for up to 60% of the proposed annotations (**Supplementary Tables S7-S8**). Most of the mistaken annotations provided by MS-DIAL came from the DIA data or from incorrect adduct annotations.

In addition, LipidMS provided a higher level of structural information for many of the annotated lipids thanks to the use of intensity ratios between characteristic fragments to elucidate the position of each fatty acyl moiety in the lipid structure (**Supplementary Tables S7-S8**). It should also be noted that while LipidMS can handle full scan, DDA and DIA data simultaneously, DIA datasets have to be processed separately for MS-DIAL.

## 3 Conclusions

Lipid annotation remains a bottleneck in lipidomic data analyses. To overcome it, LipidMS 3.0 offers an integral workflow that allows high-confidence lipid identification and quantification using DDA and DIA data and fragmentation rules. Overall, we demonstrate the significant improvements achieved by the new LipidMS release and how its use complements the information provided by existing tools. LipidMS 3.0 pre-processing capabilities are comparable to those of XCMS 3.16, and it provides complementary lipid annotation capabilities to those offered by MS-DIAL 4.80. While MS-DIAL provides wider coverage, LipidMS can simultaneously handle DDA and DIA data, provides higher structural information and returns a much smaller number of incorrect annotations (**Supplementary Figure S16**). We believe that LipidMS 3.0 is a valuable addition to existing tools and has the potential to become a key resource to annotate complex lipids. LipidMS 3.0 is an open-access platform-independent software that can be executed locally with an R environment (https://CRAN.R-project.org/package=LipidMS) or online by http://www.lipidms.com with no installation requirements.

