## Supplementary Information for "LipidMS 3.0: an R-package and a web-based tool for LC-MS/MS data processing and lipid annotation"

**Supplementary Results.**

### 1. Features and implementation

#### 1.1. Data pre-processing

Since its first version, LipidMS has been updated to cover the whole workflow required to analyze untargeted lipidomics LC-MS datasets:

***Peak-picking***. The first step of the LipidMS 3.0 workflow consists in data pre-processing using the *dataProcessing* function, which extracts all the peaks from each sample in the dataset. This function is based on the enviPick algorithm (https://CRAN.R-project.org/package=enviPick), which has been incorporated into LipidMS 3.0. At this point, $^{13}C$ isotopes are annotated. The following criteria must be met by a peak to be considered a $^{13}C$ isotope: i) mass difference of 1.0033 between the $^{12}C$ and the $^{13}C$ isotopes; ii) relative intensity between isotopes, consistently with the known natural abundances of $^{12}C$ and $^{13}C$; iii) co-elution, calculated using Pearson correlation based on peak shape as described in (Alcoriza-Balaguer *et al.*, 2019). The output is an *msobject* containing a *peaklist* and raw data for single samples. Additionally within LipidMS 3.0, the *batchdataProcessing* function can be used, which returns *msobjetcs* for each sample and wraps them into an *msbatch,* which will be subsequently used for the next batch data processing steps.

***Peak alignment***. Once all the peaks have been extracted, the time drifts during the acquisition queue need to be corrected. The *alignmsbatch* function performs peak alignment based on the MS1 information obtained for all the samples. First, peak partitions are created based on the enviPick algorithm to speed up the following clustering algorithm. Briefly, peaks are ordered by increasing mass-to-charge ratio (*m/z*) and retention time (RT), and are grouped based on user-defined tolerances. Each peak is initialized as a partition and then evaluated to decide whether it can be joined to the previous partition or not. If the *m/z* and RT of a peak match the tolerances of any of the peaks in the previous partition, this peak is reassigned. Then the clustering algorithm (**Supplementary Figure S1**) is executed to group peaks based on their RT following the next steps for each partition:

1. Each peak in the partition is initialized as a new cluster. For each cluster, the minimum, maximum and mean values of the RT which, at this point, have the same values, are kept.
2. Calculate a distance matrix between all the clusters. This distance will be the biggest difference between the minimum and maximum values of each cluster. Distances between the clusters containing peaks from the same samples will be set at NA (i.e. if a sample has two peaks in two different clusters, these two clusters cannot be merged).

3. If any distance differs from NA, search the minimum distance between two clusters.
4. If distance is below the maximum distance allowed, join clusters and update the minimum, maximum and mean values. Otherwise, set the distance to NA and go back to point 3.

Then the clusters with a sample representation over a defined minimum will be used for alignment. To this end, an RT matrix that contains the RT of the peaks for each sample from the selected clusters is built. Then the median RT is calculated for each cluster and an RT deviation matrix is obtained. Finally, the time drifts for each sample are corrected using LOESS regression by constructing a function based on RT deviation and the median. This same function is used to correct time drifts in MS2 for those samples acquired in the DIA or DDA modes.

***Peak grouping***. Once alignment has been performed, the peaks from the different samples that belong to the same metabolite/feature are grouped using the *groupmsbatch* function. To this end, the same algorithms as those employed for alignment are applied in the following order: peak partitions are created based on the *m/z* and RT values using the enviPick algorithm; *m/z* clustering is applied to each partition as described previously for RT; then peaks are grouped by RT using the same clustering algorithm. Finally, the clusters with a sample representation over a defined minimum are selected to build the feature table. An example of sequential partitioning and clustering executed for the alignment and grouping steps is summarized in **Supplementary Figure S2**.

***Filling peaks***. Once all feature peaks have been defined, areas are extracted again for each peak and sample based on the peak parameters defined for each feature (*m/z* and tolerance and initial and end RT) using the *fillpeaksmsbatch* function.

#### 1.2. Lipid annotation

Lipid annotation based on the DIA or DDA acquired samples is performed with the *annotatemsbatch* function to search for lipids in the *msbatch* based on a set of predefined fragmentation rules. As explained in the original LipidMS publication (Alcoriza-Balaguer *et al.*, 2019), only those features confirmed as M+0 (for which at least M+1 has been detected) are used for annotation to reduce false-positive annotations. Depending on the structural evidence acquired for annotation, lipids can be identified with four different confidence levels: (i) “MS-only”, when no clear fragmentation pattern is known (available only for monoacylglycerols and fatty acids); (ii) “subclass level”, when specific subclass fragments are found, but only the total number of carbons and double bonds of the chains can be proposed based on the precursor ion; (iii) “fatty acyl level”, when specific chain fragments inform about the composition of fatty acyl chains, but no positional information can be provided; (iv) “fatty acyl position level”, when the specific chain position can be elucidated based on chain fragments’ intensity ratios. Ever since LipidMS was first released, six new lipid classes have been included to, thus, cover 30 lipid classes. The adducts and fragmentation rules for these new classes, based on experimental spectra and publicly available spectra databases, are summarized in **Supplementary Tables S1-S3**. The experimental data supporting these rules are represented in **Supplementary Figures S3-S14**.

#### 1.3. LipidMS workflow

LipidMS supports the simultaneous processing of all the possible combinations of MS acquisitions modes:

- All the samples in DIA
- All the samples in DDA
- Combination of DDA and DIA
- Combination of full scan and DIA
- Combination of full scan and DDA
- Combination of full scan, DIA and DDA

As specified in Figure 1, the workflow is:

i. Data input

Load mzXML files and associated metadata in the .csv format

ii. Data pre-processing

Peak picking is performed for MS1 in all the samples, and MS2 in those files acquired in the DIA mode. Then grouping and alignment are performed based on the information contained in MS1. For those samples acquired in DIA or DDA, MS2 scans are aligned based on the parameters obtained for MS1.

For DIA, the link between MS1 and MS2 is performed based on RT windows and the peak shape correlation as specified in the original manuscript (Alcoriza-Balaguer *et al.*, 2019).

For DDA, the link between MS1 and MS2 is based on RT differences. For each feature of interest (MS1 peak that potentially matches a lipid), MS2 data are searched. To this end, the algorithm searches for the MS2 scans that fall within the limits of the MS1 peak. If multiple MS2 scans meet requirements, the average spectra are used. If an MS2 scan falls within the limits of two MS1 peaks, i.e. closely eluting isomeric or isobaric species, the MS2 scan is assigned to the closest MS1 peak.

The result of step 2 is a feature matrix that contains information for all the detected features across all the samples.

iii. Lipid annotation

Lipid annotation is performed based on fragmentation rules as specified in the original paper (Alcoriza-Balaguer *et al.*, 2019) and in **Supplementary Tables S1-S3** for each sample acquired in DIA or DDA. First, putative annotations are performed based on the *m/z* and features corresponding to the same assigned lipid (e.g. $[M+H]^+$ and $[M+Na]^+$ adducts). Then fragmentation rules are employed to perform the actual lipid annotation.

If different annotation levels are obtained for a given lipid, e.g. feature X is identified as PC(34:1) based on DIA and PC(16:1/18:0) based on DDA, the identification with the highest degree of structural information [i.e. PC(16:1/18:0)] is maintained. If several identifications with the same annotation level are obtained [e.g. PC(18:1/16:0), PC(16:1/18:0)], they are all maintained.

Finally, lipid annotations are incorporated into the feature matrix.

iv. Output

The final outputs are a table containing the signal for each identified lipid across all the samples and a pdf with the data supporting each identification.

#### 1.4. Web application

In order to provide a user-friendly GUI interface, LipidMS 3.0 has also been implemented as a web-based tool, which is accessed at http://www.lipidms.es. After accessing the tool, the following tabs will take users through the LipidMS workflow:

***Data import.*** In the first tab, users must choose polarity and upload all the mzXML files and a metadata file in the csv format with three columns: sample (mzXML file names); acquisition mode (MS for full scan, DIA or DDA); sample type (QC, group1, group2, etc.).

***Peak-picking.*** Then all the parameters required for peak-picking can be tuned. In this tab, the MS1 and MS2 values correspond to those parameters used to process the MS1 level in all cases and the MS2 level for the DIA data files, respectively.

***Batch processing.*** The third tab contains the parameters required for alignment, grouping and filling peak steps.

***Annotation.*** In the annotation tab, the lipid classes to be searched, and the *m/z* and RT tolerances, can be defined.

***Run.*** Finally, users can run their job. The results will be sent to the provided email containing two or three types of csv files with the results tables (feature matrix if batch processing is performed, summary tables and the whole peak tables with annotations) and the pdf files with plots of the peaks supporting the lipid identifications for all the files.

Extra documentation and examples and links to the source code in github or CRAN can be found by clicking on the "Resources" tab. Web tool screenshots are shown in **Supplementary Figure S15**.

### 2. LipidMS 3.0 performance evaluation

#### 2.1. Identification of lipid standards spiked into a human serum pool

A commercial human serum pool (Sigma-Aldrich reference P2918) was analyzed by LC-MS using both the ESI + and ESI ionization modes and in full scan, DIA and DDA acquisition modes. To provide an objective qualifier for the comparison, serum was extracted with or without the addition of 68 lipid standards. Raw data files and results are available at Zenodo (https://doi.org/10.5281/zenodo.6645498).

Three different workflows were compared: 1) LipidMS workflow; 2) MS-DIAL workflow (Tsugawa *et al.*, 2015); 3) data pre-processing using XCMS (Smith *et al.*, 2006), isotope annotation using CAMERA (Kuhl *et al.*, 2012).

When employing the LipidMS workflow (i.e., data pre-processing and lipid annotation), the samples acquired in full scan, DIA and DDA were simultaneously processed, which is not possible with MS-DIAL. For MS-DIAL, the MS and DDA files were processed together and the DIA files separately. Then DIA identifications were added to the feature/identities matrix obtained for the full scan and DDA files using a *m/z* tolerance of 0.005 and an RT tolerance of 10 seconds. XCMS was exclusively applied to pre-process the MS1 level of all the samples and CAMERA for the annotation of the isotopes in the generated feature matrix. In all cases, the reported features referred to the combination of the positive and negative ionization modes for the MS1 level, and lipid identities are based on the information obtained for MS2 from the DDA and DIA acquired samples. In all three cases, features were filtered and normalized based on quality control (QC) samples. Only the features that appeared in at least 70% of the QC samples were kept. Then data were normalized using a LOESS function, which was fitted to the QC samples based on the injection order. The resulting interpolated curve for each feature was used to normalize its response (Dunn *et al.*, 2011). Finally, a differential analysis between the spiked and non spiked serum samples was performed using a Student's t-test, corrected for multiple testing and magnitude of change. The features with an adjusted *p-value* < 0.05 and a fold change > 1.5 were considered to be differential variables between both groups.

The performance of each pre-processing software was evaluated by comparing the number of detected lipid standard features (a single lipid can be annotated using different features that could result from the ionization of many adducts, e.g. $[M+H]^+$ and $[M+Na]^+$, thus, 129 features were expected from the 68 lipids) and the differences in lipid standard features between spiked and non spiked samples. Additionally, for the comparison between MS-DIAL and LipidMS workflows the number of correctly annotated lipid standard features and lipid standards were evaluated.

Comparison between LipidMS and XCMS data pre-processing

Previous LipidMS versions were designed for lipid annotation of single samples, which required the use of external software like XCMS to perform the LC-MS untargeted analysis. Now LipidMS 3.0 includes the whole workflow for batch processing (i.e., peak-peaking, alignment and grouping). To evaluate the performance of these pre-processing steps incorporated into LipidMS 3.0, we compared the results obtained by the LipidMS 3.0 workflow to those obtained by employing one of the most widely used pre-processing platforms: XCMS (Smith *et al.*, 2006).

Despite the fact that XCMS 3.16 found a larger total number of features than LipidMS 3.0 (33352 in XCMS vs. 19382 in LipidMS) (**Supplementary Table S4**), both software provide similar numbers in terms of the expected lipid standard features detected and significant changes in them (**Supplementary Tables S4-S6**). These results validate the LipidMS pre-processing workflow, which for the lipidomic studies provided results that were comparable to those obtained by XCMS 3.16.

Comparison between LipidMS and MS-DIAL

MS-DIAL found a larger number of features than LipidMS (75263 vs. 25574) (**Supplementary Table S4**), but MS-DIAL and LipidMS provided similar numbers for the expected lipid standard features detected and significant changes in them (**Supplementary Tables S4-S6**). However for the lipid standards, LipidMS provided a larger number of both identified features (98/129 vs. 76/129) and identified lipids (60/68 vs. 56/68) (**Supplementary Tables S4-S6**). Most of the differences in the proposed identities came from the MS-DIAL incorrect assignation of some adducts, where ions $[M+H]^+$ and $[M-H]^-$ were correctly annotated, but adducts like $[M+Na]^+$, $[M+CH_3COO]^-$, $[M-CH_3]^-$ or $[M+Na-2H]^-$ were not annotated or incorrectly annotated (**Supplementary Tables S5-S6**). Thus by means of this strategy, which focused on a subset of lipid classes covered by MS-DIAL and LipidMS, both software packages provided comparable results or LipidMS even slightly outperformed MS-DIAL. The improved LipidMS annotation of adducts compared to MS-DIAL was due to its underlying lipid annotation strategy in which features were first assigned as being related (e.g. putative $[M+H]^+$ and $[M+Na]^+$ ions of a given lipid), and then the lipid identity was proposed. This approach reduces the possibility of proposing different lipid identities for adduct features from a single lipid.

### 2.2. Annotation of lipids in a human serum pool sample

LipidMS performance was compared to MS-DIAL in relation to the total number of lipid annotations provided for the aforementioned commercial human serum pool (Sigma-Aldrich reference P2918) analyzed by LC-MS in both the ESI + and ESI – ionization modes and by full scan, DIA and DDA acquisition methods. All the annotations provided by MS-DIAL and LipidMS 3.0 were manually curated and the results compared. The raw data files and an Excel file containing the curated lipid identities, the annotations proposed by LipidMS and the annotations proposed by MS-DIAL can be accessed at Zenodo (https://doi.org/10.5281/zenodo.6645498).

For both polarities, MS-DIAL provided a larger number of correct lipid annotations (580 vs. 387 in ESI – and 588 vs. 445 in ESI+) (**Supplementary Tables S7-S8**). The main reasons for the increased coverage provided by MS-DIAL were: 1) more lipid classes covered by MS-DIAL; 2) higher diversity of fatty acyl chains in terms of chain length and double bonds, and also due to the inclusion of oxidized and hydroxylated fatty acyl chains; 3) higher diversity of ceramides and sphingomyelins. Based on the results of this comparison, LipidMS will incorporate new lipid classes, sphingoid bases and fatty acyl moieties to bridge the gap in lipid coverage with MS-DIAL. Conversely, LipidMS provided higher structural information compared to MS-DIAL (i.e. a bigger proportion of lipids where the structural information level achieves an FA position). LipidMS uses the ratio between the intensity of fragments to elucidate the position of fatty acyl chains for most lipid classes, whereas MS-DIAL only discloses the fatty acyl position for ceramides and sphingomyelins. Finally in both polarities, MS-DIAL also provided more incorrect annotations (669 vs. 79 in ESI – and 897 vs. 50 in ESI+). For LipidMS, incorrect annotations represented below 20% of the proposed annotations, but added up to 60% of the proposed identities for MS-DIAL (**Supplementary Tables S7-S8**). Most incorrect annotations came from the DIA data, where many annotations were based on noisy spectra and the majority of the reference ions were not present in the samples. As previously mentioned, many incorrect annotations were due to the incorrect assignment of adducts. This was particularly relevant for cardiolipins because almost all of them were incorrectly annotated, and some phosphatidylcholines and phosphatidylethanolamines were erroneously assigned to a particular class due to the miss-annotation of $[M+Na]^+$ as $[M+H]^+$ or $[M+CH3COO]^-$ as $[M-H]^-$.

In short, when the performances of LipidMS 3.0 and MS-DIAL to annotate lipids in a complex biological sample were compared, MS-DIAL annotated more lipids, but also provided more incorrect annotations, peaks that did not correspond to known lipids and were annotated or lipids with a miss-annotation, whereas

LipidMS provided fewer annotated lipids, as well as fewer false-positives and higher level of structural information (**Supplementary Tables S7-S8, Supplementary Figure S16**). Although MS-DIAL separately processes DDA and DIA files, any combination can be simultaneously processed using LipidMS

**Supplementary Methods.**

***Chemicals.*** The solvents and additives for sample processing and LC-MS analysis were isopropanol, formic acid, ammonium formate and ammonium acetate, obtained from Sigma-Aldrich/Fluka (Madrid, Spain), and acetonitrile supplied by Fisher Scientific (Loughborough. UK).

Lipid standards were obtained from Avanti Polar Lipids (Alabaster, AL, USA), Sigma-Aldrich/Fluka (Madrid, Spain) and Larodan (Solna, Sweden). Lipid standards were 1-O-oleoyl-N-heptadecanoyl-D-sphingosine [AcylCer(18:1;18:1/17:0)], cholest-5-en-3b-yl heptadecanoate [CE(17:0)], N-heptadecanoyl-D-sphingosine [Cer(d18:1/17:0)], N-palmitoyl-D-sphingosine-1-phosphate [CerP(d18:1/16:0)], 1,3-bis-[1,2-di-octadecenoyl-sn-glycero-3-phospho]-sn-glycerol [CL(18:1/18:1/18:1/18:1)], diheptadecanoylglycerol [DG(17:0/17:0)], capric acid [FA(10:0)], lauric acid [FA(12:0)], myristic acid [FA(14:0)], myristoleic acid [FA(14:1n5)], pentadecanoic acid [FA(15:0)], palmitic acid [FA(16:0)], palmitoleic acid [FA(16:1)n7], sapienic acid [FA(16:1n10)], margaric acid [FA(17:0)], stearic acid [FA(18:0)], vaccenic acid [FA(18:1n7t)], oleic acid [FA(18:1n9)], linoleic acid [FA(18:2n6)], alpha-linolenic acid [FA(18:3n3)], gamma-linolenic acid [FA(18:3)n6], nonadecanoic acid [FA(19:0)], arachidic acid [FA(20:0)], gondoic acid [FA(20:1n9)], 11,14-eicosadienoic acid [FA(20:2n6)], dihomo-alpha-linolenic acid [FA(20:3n3)], dihomo-gamma-linolenic acid [FA(20:3n6)], 8,11,14,17-eicosatetraenoic acid [FA(20:4n3)], arachidonic acid [FA(20:4n6)], eicosapentaenoic acid [FA(20:5n3)], behenic acid [FA(22:0)], erucic acid [FA(22:1n9)], docosadienoic acid [FA(22:2n6)], 10,13,16-docosatrienoic acid [FA(22:3n6)], 10,13,16,19-docosatetraenoic acid [FA(22:4n3)], adrenic acid [FA(22:4n6)], clupanodonic acid [FA(22:5n3)], 4,7,10,13,16-docosapentaenoic acid [FA(22:5n6)], cervonic acid [FA(22:6n3)], lignoceric acid [FA(24:0)], nervonic acid [FA(24:1n9)], cerotic acid [FA(26:0)], heptadecanoyl-sn-glycero-3-phosphocholine [LPC(17:0)], monoheptadecanoylglycerol [MG(17:0)], 1-hexadecanoyl-2-octadecenoyl-sn-glycero-3-phosphocholine [PC(16:0/18:1)], 1,2-diheptadecanoyl-sn-glycero-3-phosphatidylcholine [PC(17:0/17:0)], 1-octadecanoyl-2-octadecadienoyl-sn-glycero-3-phosphocholine, [PC(18:0/18:2)], 1-hexadecyl-2-(5Z,8Z,11Z,14Z,17Z-eicosapentaenoyl)-sn-glycero-3-phosphocholine, [PC(O-16:0/20:5)], 1-(1Z-octadecenyl)-2-(5Z,8Z,11Z,14Z-eicosatetraenoyl)-sn-glycero-3-phosphocholine [PC(P-18:0/20:4)], 1,2-diheptadecanoyl-sn-glycero-3-phospholethanolamine [PE(17:0/17:0)], 1-hexadecanoyl-2-octadecenoyl-sn-glycero-3-phosphoetyhanolamine [PE(16:0/18:1)], 1-hexadecyl-2-(9Z-octadecenoyl)-sn-glycero-3-phosphoethanolamine [PE(O-16:0/18:1)], 1-(1Z-octadecenyl)-2-(4Z,7Z,10Z,13Z,16Z,19Z-docosahexaenoyl)-sn-glycero-3-phosphoethanolamine [PE(P-18:0/22:6)], 1-hexadecanoyl-2-octadecenoyl-sn-glycero-3-phosphoglycerol [PG(16:0/18:1)], 1,2-diheptadecanoyl-sn-glycero-3-phosphoglycerol [PG(17:0/17:0)], 1-heptadecanoyl-2-(9Z-tetradecenoyl)-sn-glycero-3-phospho-(1'-myo-inositol) [PI(17:0/14:1)], 1-hexadecanoyl-2-octadecenoyl-sn-glycero-3-phosphoserine [PS(16:0/18:1)],

1,2-diheptadecanoyl-sn-glycero-3-phosphoserine [PS(17:0/17:0)], N-palmitoyl-D-sphingomyelin [SM(18:1/16:0)], N-heptadecanoyl-D-sphingomyelin [SM(18:1/17:0)], 1,2,3-octanoylglycerol [TG(8:0/8:0/8:0)], 1.2.3-tridecanoylglycerol [TG(10:0/10:0/10:0)], 1,2,3-tridodecanoylglycerol [TG(12:0/12:0/12:0)], 1,2,3-tritetradecanoylglycerol [TG(14:0/14:0/14:0)], 1,2,3-trihexadecanoylglycerol [TG(16:0/16:0/16:0)], 1,2-dipalmitoyl-3-oleoylglycerol [TG(16:0/16:0/18:1)], 1,2,3-triheptadecanoylglycerol [TG(17:0/17:0/17:0)], 1.3-dioleoyl-2-palmitoylglycerol [TG(18:1/16:0/18:1)].

***Sample preparation.*** For lipid extraction, 50 µL of human serum were mixed with 10 µL of solvent or a mixture of lipid standards at 20 µg/mL each and 150 µL of isopropanol. After vortexing, samples were left for 20 min at -20ºC and then centrifuged for 15 min at 15000g and 4ºC. Finally, 100 µL of the supernatants were transferred to an HPLC vial for their LC-MS-based analysis.

***UPLC-HRMS analysis.*** The lipidomic profiles of human serum samples were analyzed by a quadrupole–orbitrap mass spectrometer (Q Exactive, Thermo Fisher Scientific) coupled to reverse phase chromatography via electrospray ionization. Liquid chromatography separation took place in a CSH C18 column (2.1 mm × 100 mm, 1.7 µm particle size; Waters). Solvent A was 10 mM ammonium acetate (ESI +) or ammonium formiate (ESI -) in 60:40 acetronitile:water. Solvent B was 10 mM ammonium acetate (ESI +) or ammonium formiate (ESI -) in 90:10 isopropanol:acetonitrile. The flow rate was 0.4 µL/min, the column temperature was 65 °C, the autosampler temperature was 5 °C and the injection volume was 5 µL. The liquid chromatography gradient was as previously described (Alcoriza-Balaguer *et al.*, 2019). Briefly, 0 min 20% B, 2 min 20% B, 4 min 43% B, 4.10 min 50% B, 14 min 54% B, 14.10 min 70% B, 20 min 99% B, 24 min 99% B, 24.50 min 20% B and 27 min 20% B at a flow of 0.4 mL/min. The following conditions were employed for the ESI+ and ESI- ionization modes, respectively: the sheath gas flow rate was 25 (0-14 min) and 80 (14-27 min) for ESI+ and 25 for ESI-; the auxiliary gas flow rate was 10 (0-14 min) and 25 (14-27 min) for ESI+ and 25 for ESI-; the spray voltage was 3 kV for ESI+ and 2.5 kV for ESI-; the capillary temperature was 215 ºC for ESI+ and 400 ºC for ESI-; the S-lens RF-level was 95 for ESI+ and 65 for ESI-; the auxiliary gas heater temperature was 215ºC for ESI+ and 350ºC for ESI-. The pooled samples were acquired in full scan, DDA and DIA modes, while individual samples were acquired only in the MS scan mode. For MS scan acquisition purposes, resolution was set at 70000, the AGC target at 1000000, the maximum IT at 100 ms, the scan range was 113-1700 and data type was centroid. For DDA acquisition purposes, the full scan parameters were as in the MS acquisition, while the MS2 parameters were: resolution 70000, AGC target 1000000, maximum IT 200 ms, loop count 5, MSX count 1, isolation window 0.4 *m/z*, isolation offset 0.4 *m/z*, collision energies 30 and 40 V, data type centroid, minimum AGC target 1000 and dynamic exclusion 5 sec. Finally for DIA acquisition purposes, the full scan parameters were as in the MS acquisition, while the MS2 parameters were: resolution 70000, AGC target 1000000, maximum IT 200 ms, collision energies 30 and 40 V, scan range 80-1200 and data type centroid.

***Data processing parameters.***

- LipidMS 3.0: all the samples were processed together (full scan, DDA and DIA) using the LipidMS R package. Files were previously converted into the mzXML format using the msConvert software (ProteoWizard 3.0.10800).
    - Peak-picking parameters:
        - dmzagglom: 15
        - drtagglom: 200
        - drtclust: 25
        - minpeak: 5
        - drtgap: 5
        - drtminpeak: 10
        - drtmaxpeak: 200
        - recurs: 5 for MS1 and 10 for MS2.
        - sb: 3 for MS1 and 2 for MS2.
        - sn: 3 for MS1 and 2 for MS2.
        - minint: 1000 for MS1 and 100 for MS2.
        - weight: 2 for MS1 and 3 for MS2.
        - dmzIso: 5
        - drtIso: 5
    - Batch processing parameters (alignment and grouping):
        - dmzalign: 10
        - drtalign: 100
        - span: 0.4
        - minsamplesfracalign: 0.75
        - dmzgroup: 10
        - drtagglomgroup: 50
        - drtgroup: 15
        - minsamplesfracgroup: 0.30
    - Lipid annotation parameters:
        - dmz for precursors: 5,
        - dmz for products: 10

        - rttol: 6,
        - coelCutoff: 0.6

- XCMS 3.16: all the samples were pre-processed together for MS level 1 (full scan, DDA and DIA) by the XCMS R package. Files were previously converted into the mzXML format using the msConvert software (ProteoWizard 3.0.10800). Several values for the bandwidth and binSize parameters were tested to optimize the extraction of isomeric peaks.
    - Peak-picking:
        - peakwidth: between 5 and 30 seconds
        - noise: 1000
        - ppm: 15
        - snthres: 3
        - prefilter: 5 scans with a minimum intensity of 1000
    - Alignment and grouping:
        - Alignment method: based on the peak groups.
        - Grouping:
            - minFraction: 0.3
            - bw = 2
            - binSize = 0.005

- CAMERA 3.15: The XCMS pre-processed files were processed using CAMERA 3.15 for the annotation of isotopes.
        - perfwhm = 0.6
        - cor_eic_th = 0.75
        - maxcharge = 3
        - ppm = 5
        - mzabs = 0.01
        - filter (C12/C13) = TRUE

- MS-DIAL 4.80: the full scan and DDA acquired samples were processed together, while the DIA files were analyzed in a different batch. Files were previously converted into the abf format using Reifycs Abf (Analysis Base File) Converter 4.0. Then the DIA identifications were added to the feature matrix obtained for the full scan and DDA files using an *m/z* tolerance of 0.005 and an RT tolerance of 10 seconds.
    - Data collection:

        - MS1 tolerance: 0.005
        - MS2 tolerance: 0.01
    - Peak detection:
        - Minimum peak height: 1000
        - Mass slice width : 0.1 Da
        - Smoothing method: Linear-weighted moving average
        - Smoothing level 5 scan
        - Minimum peak width: 5 scan
    - Adducts:
        - ESI-: M-H, M-H-H2O, M+Na-2H, M+Hac-H, M+FA-H, 2M-H and M-2H.
        - ESI+: M+H, M+NH4, M+Na, M+H-H2O, M+H-2H2O, 2M+NH4 and 2M+Na.

**Supplementary References.**

Alcoriza-Balaguer,M.I. *et al.* (2019) LipidMS: An R Package for Lipid Annotation in Untargeted Liquid Chromatography-Data Independent Acquisition-Mass Spectrometry Lipidomics. *Anal. Chem.*, **91**, 836–845.

Dunn,W.B. *et al.* (2011) Procedures for large-scale metabolic profiling of serum and plasma using gas chromatography and liquid chromatography coupled to mass spectrometry. *Nat. Protoc.*, **6**, 1060–1083.

Kuhl,C. *et al.* (2012) CAMERA: An integrated strategy for compound spectra extraction and annotation of liquid chromatography/mass spectrometry data sets. *Anal. Chem.*, **84**, 283–289.

Smith,C.A. *et al.* (2006) XCMS: Processing mass spectrometry data for metabolite profiling using nonlinear peak alignment, matching, and identification. *Anal. Chem.*, **78**, 779–787.

Tsugawa,H. *et al.* (2015) MS-DIAL: Data-independent MS/MS deconvolution for comprehensive metabolome analysis. *Nat. Methods*, **12**, 523–526.

### Supplementary Figures

**Supplementary Figure S1. Scheme of the clustering algorithm used for alignment and grouping**

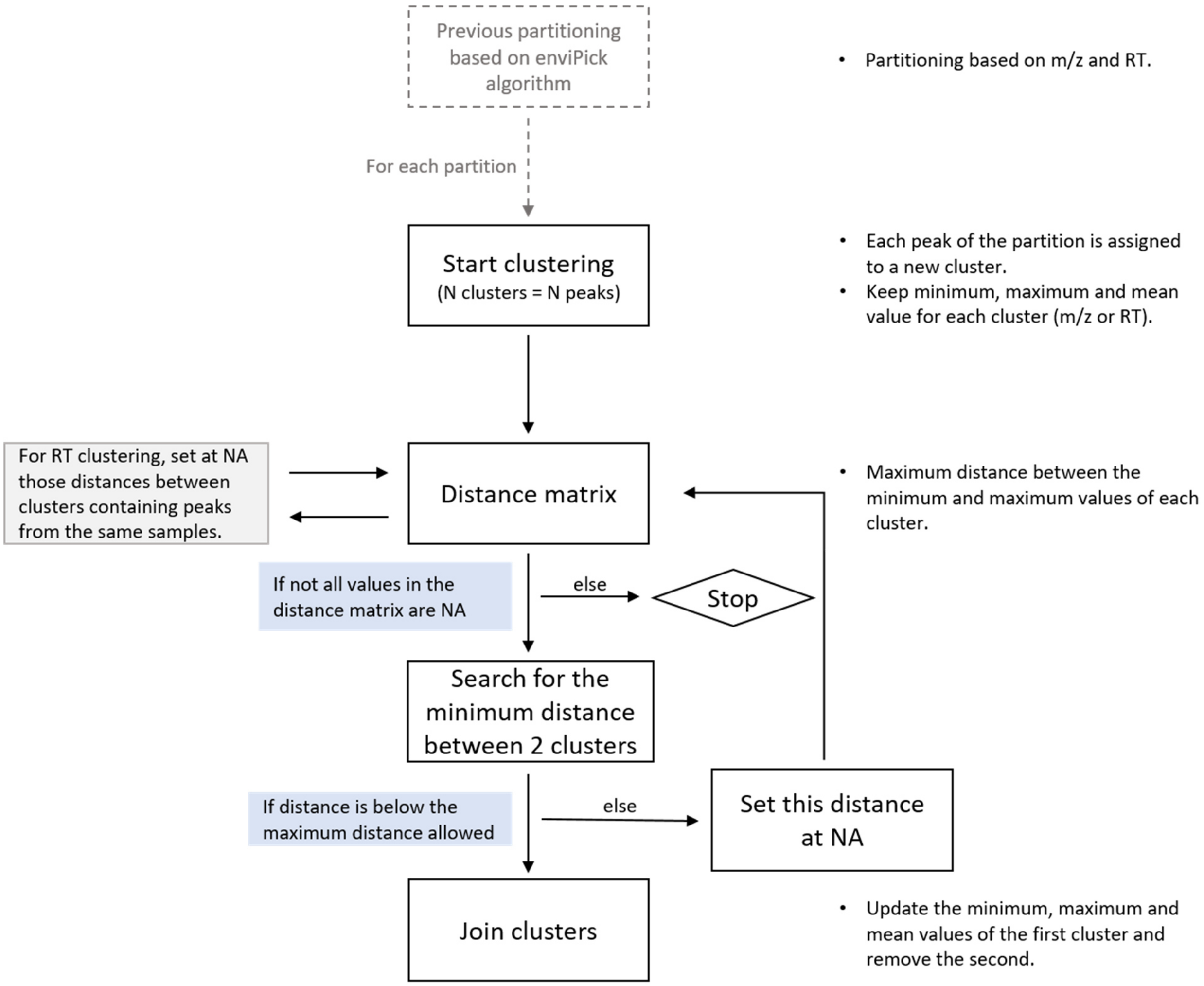

**Supplementary Figure S2. Example of the sequential partitioning and clustering of the peaks executed for the alignment and grouping steps.** Each point represents a peak, each shape denotes a sample and each color depicts a cluster.

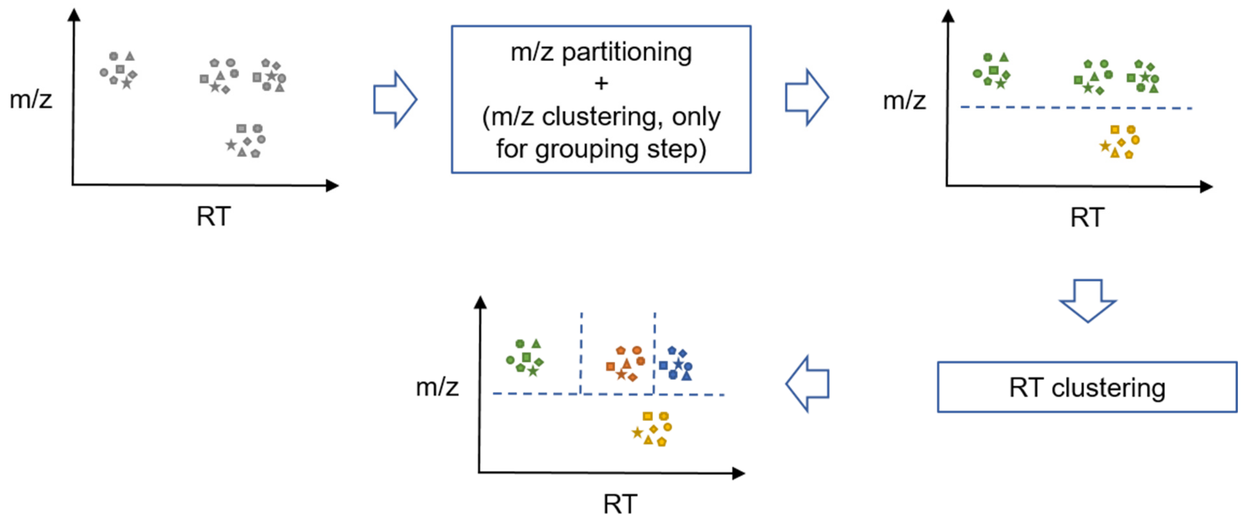

**Supplementary Figure S3. The AcylCer(18:1;d18:1/17:0) fragmentation profile for ESI+.** A) Chromatographic profiles of the precursor ions in MS1. B) Chromatographic profiles of the informative fragments in MS2 using DIA (35eV). Co-elution between the precursor and product fragments is observed (A-B). C) Fragmentation spectra for MS2 using DDA (35eV) for precursor ion 816.7808 (M+H). Only the specific fragments used for annotation are shown.

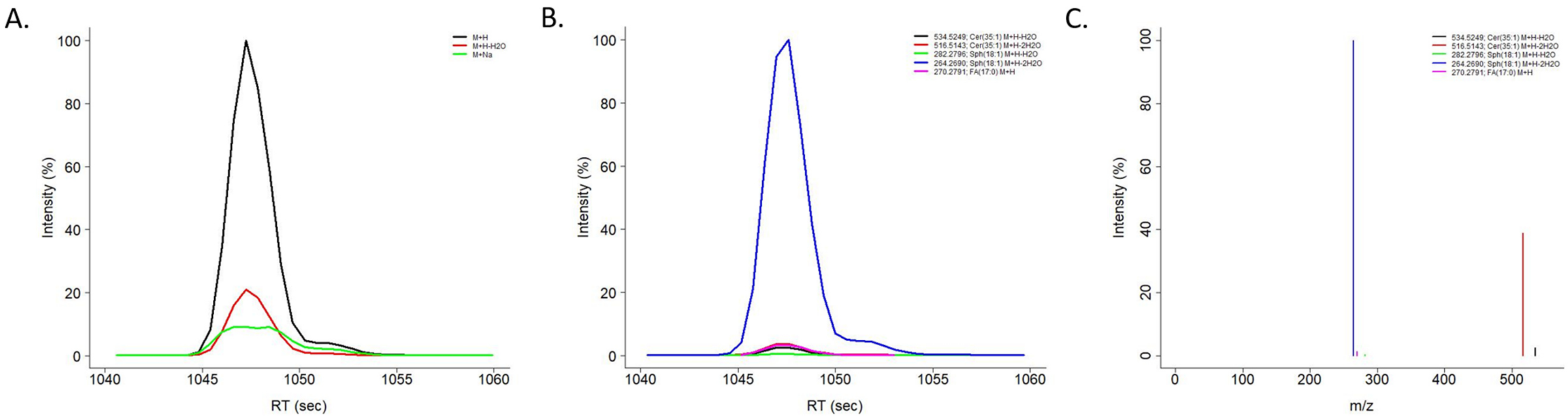

**Supplementary Figure S4. The AcylCer(18:1;d18:1/17:0) fragmentation profile for ESI-.** A) Chromatographic profiles of the precursor ions in MS1. B) Chromatographic profiles of the informative fragments in MS2 using DIA (35eV). Co-elution between the precursor and product fragments is observed (A-B). C) Fragmentation spectra for MS2 using DDA (35eV) for precursor ion 814.7652 (M-H). Only the specific fragments used for annotation are shown.

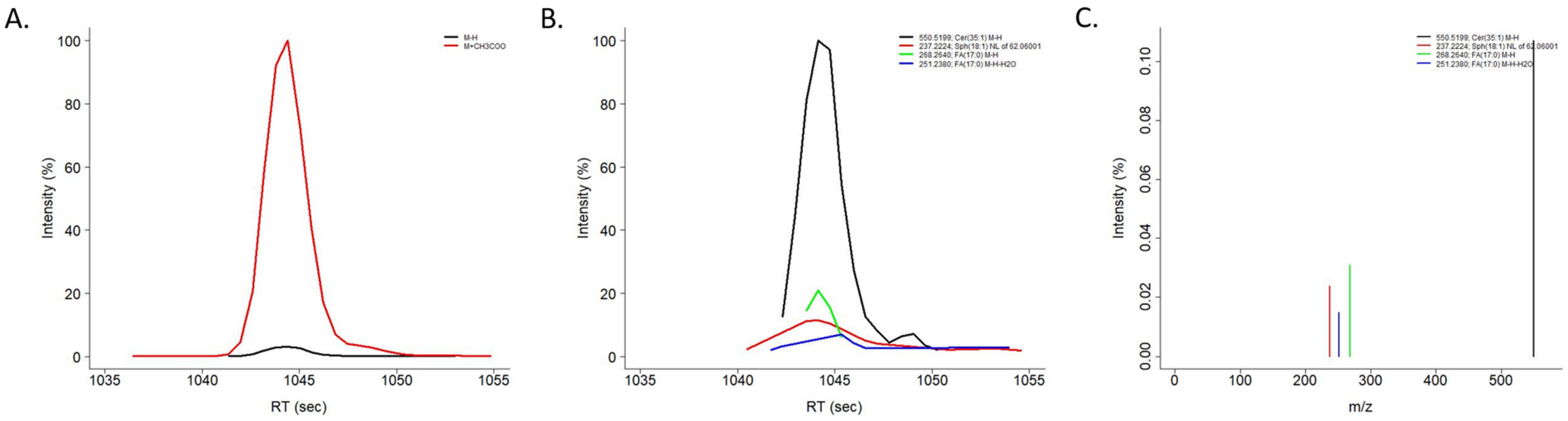

**Supplementary Figure S5. The CerP(d18:1/16:0) fragmentation profile for ESI+.** A) Chromatographic profiles of the precursor ions in MS1. B) Chromatographic profiles of the informative fragments in MS2 using DIA (35eV). Co-elution between the precursor and product fragments is observed (A-B). C) Fragmentation spectra for MS2 using DDA (35eV) for precursor ion 618.4863 (M+H). Only the specific fragments used for annotation are shown.

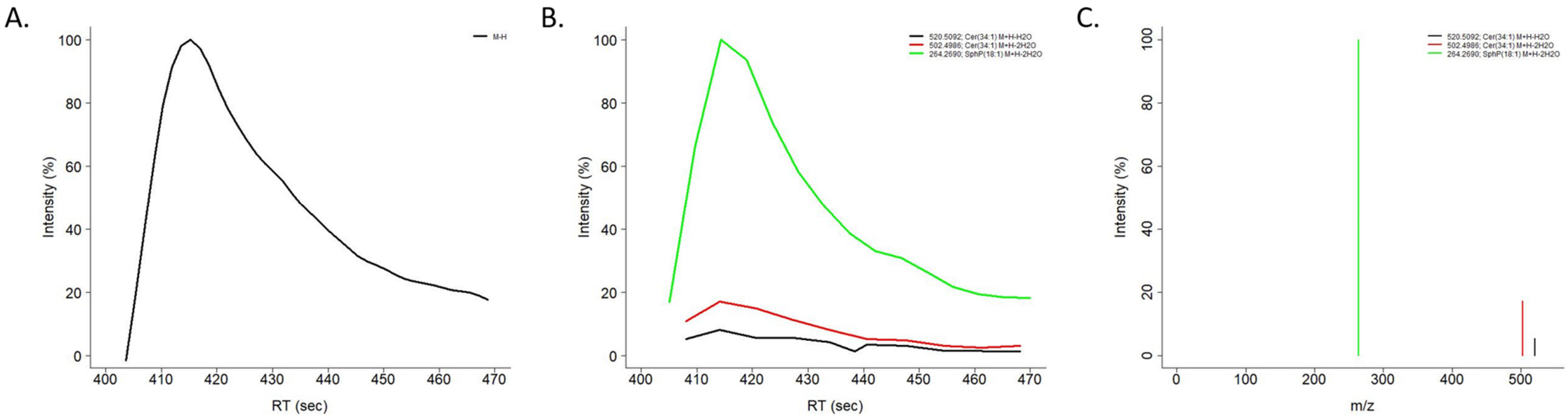

**Supplementary Figure S6. The CerP(d18:1/16:0) fragmentation profile for ESI-.** A) Chromatographic profiles of the precursor ions in MS1. B) Chromatographic profiles of the informative fragments in MS2 using DIA (35eV). Co-elution between the precursor and product fragments is observed (A-B). C) Fragmentation spectra for MS2 using DDA (35eV) for precursor ion 616.4717 (M-H). Only the specific fragments used for annotation are shown.

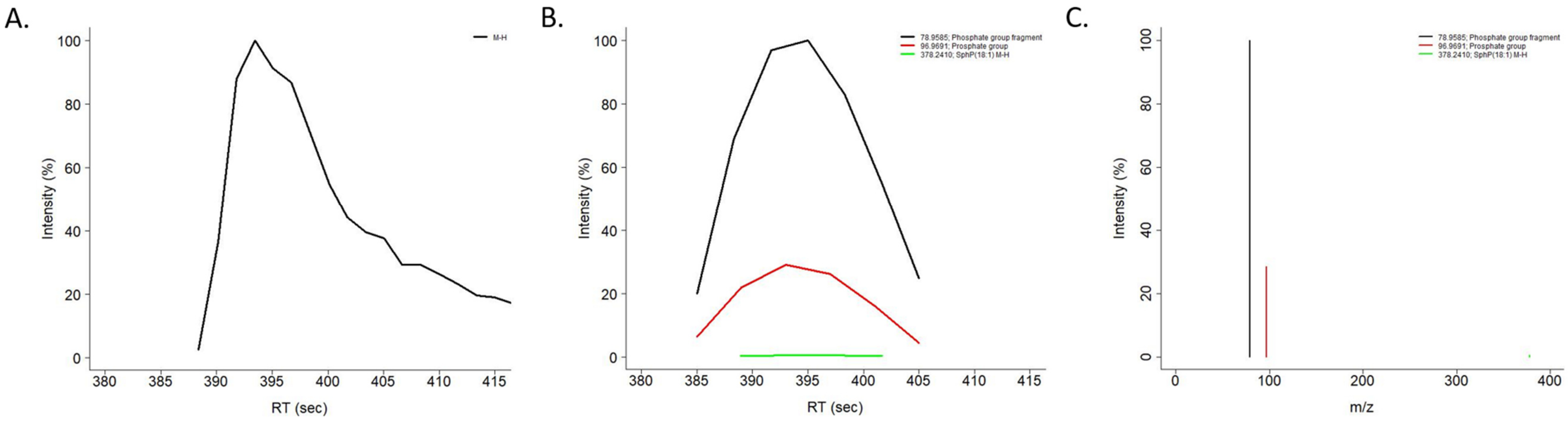

**Supplementary Figure S7. The PC(O-16:0/20:5) fragmentation profile for ESI+.** A) Chromatographic profiles of the precursor ions in MS1. B) Chromatographic profiles of the informative fragments in MS2 using DIA (35eV). Co-elution between the precursor and product fragments is observed (A-B). C) Fragmentation spectra for MS2 using DDA (35eV) for precursor ion 766.5751 (M+H). Only the specific fragments used for annotation are shown. Fragment 184 represents 100% relative intensity.

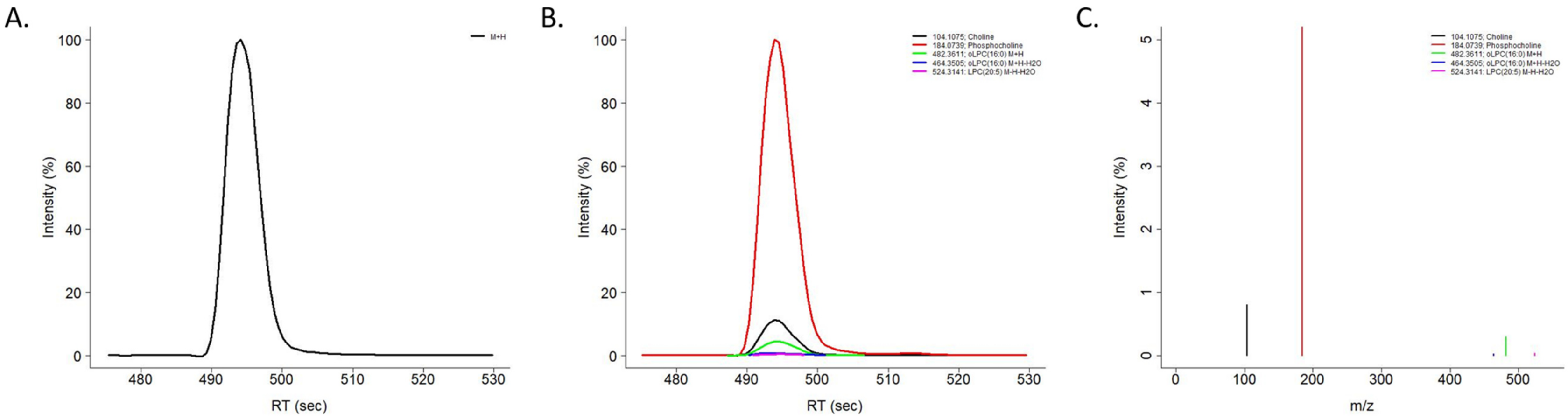

**Supplementary Figure S8. The PC(O-16:0/20:5) fragmentation profile for ESI-**A) Chromatographic profiles of the precursor ions in MS1. B) Chromatographic profiles of the informative fragments in MS2 using DIA (35eV). Co-elution between the precursor and product fragments is observed (A-B). C) Fragmentation spectra for MS2 using DDA (35eV) for precursor ion 824.5812 (M+$CH_3COO$). Only the specific fragments used for annotation are shown.

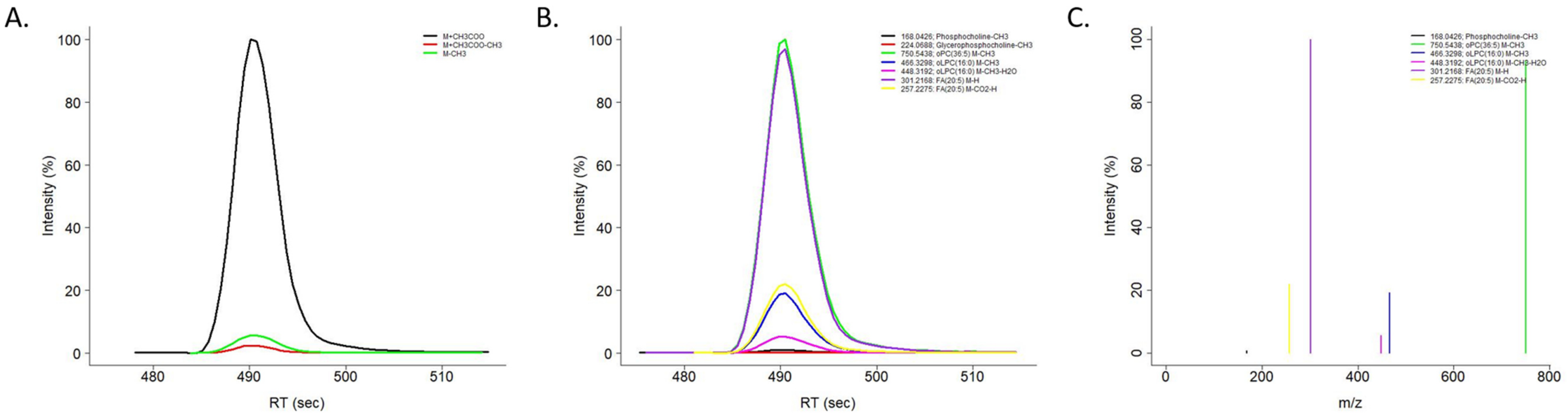

**Supplementary Figure S9. The PC(P-18:0/20:4) fragmentation profile for ESI+.** A) Chromatographic profiles of the precursor ions in MS1. B) Chromatographic profiles of the informative fragments in MS2 using DIA (35eV). Co-elution between the precursor and product fragments is observed (A-B). C) Fragmentation spectra for MS2 using DDA (35eV) for precursor ion 794.6064 (M+H). Only the specific fragments used for annotation are shown. Fragment 184 represents 100% relative intensity.

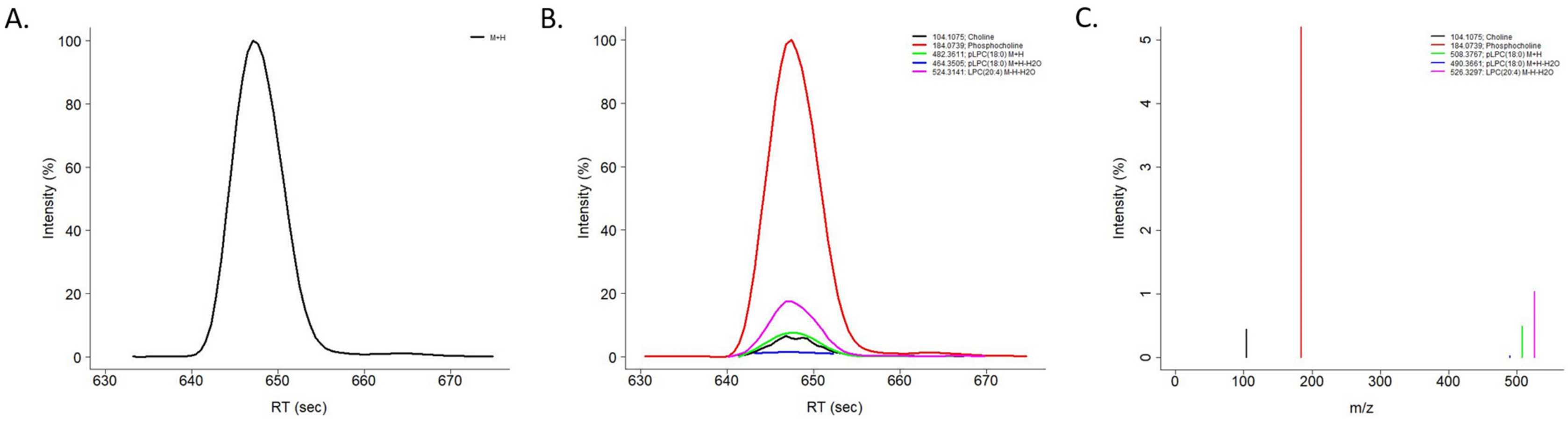

**Supplementary Figure S10. The PC(P-18:0/20:4) fragmentation profile for ESI-.** A) Chromatographic profiles of the precursor ions in MS1. B) Chromatographic profiles of the informative fragments in MS2 using DIA (35eV). Co-elution between the precursor and product fragments is observed (A-B). C) Fragmentation spectra for MS2 using DDA (35eV) for precursor ion 852.6125 (M+$CH_3COO$). Only the specific fragments used for annotation are shown.

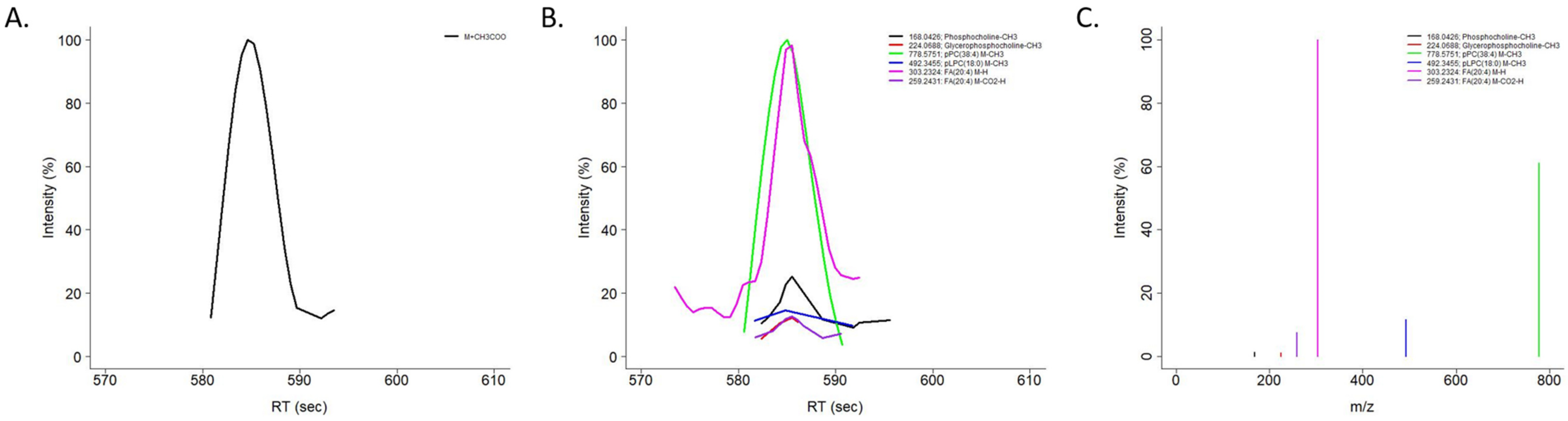

**Supplementary Figure S11. The PE(O-16:0/18:1) fragmentation profile for ESI+.** A) Chromatographic profiles of the precursor ions in MS1. B) Chromatographic profiles of the informative fragments in MS2 using DIA (35eV). Co-elution between the precursor and product fragments is observed (A-B). C) Fragmentation spectra for MS2 using DDA (35eV) for precursor ion 704.5594 (M+H). Only the specific fragments used for annotation are shown.

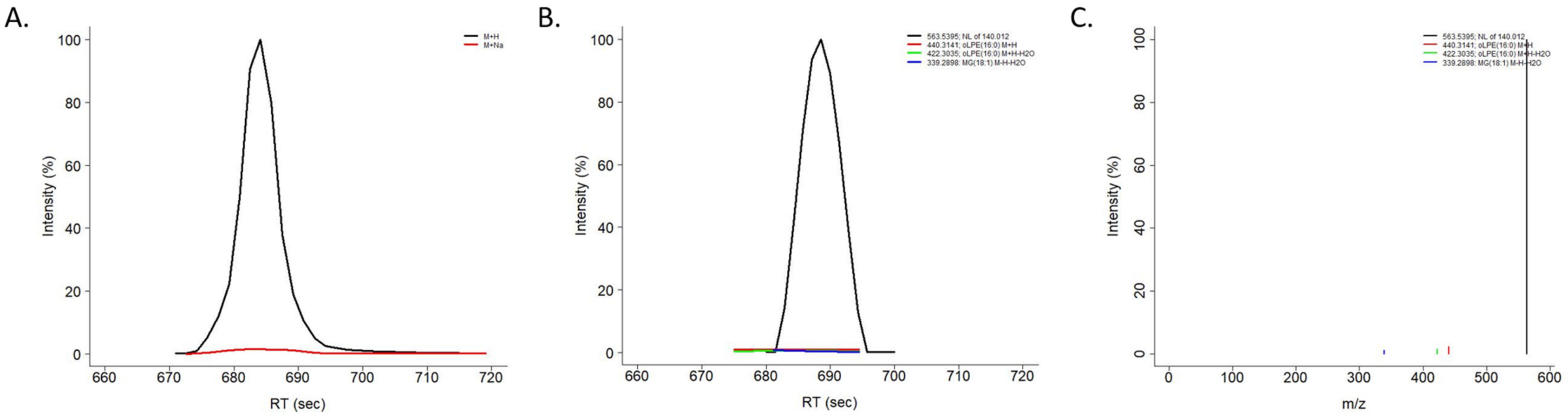

**Supplementary Figure S12. The PE(O-16:0/18:1) fragmentation profile for ESI-.** A) Chromatographic profiles of the precursor ions in MS1. B) Chromatographic profiles of the informative fragments in MS2 using DIA (35eV). Co-elution between the precursor and product fragments is observed (A-B). C) Fragmentation spectra for MS2 using DDA (35eV) for precursor ion 784.5469 (M+NaCH$_3$COO). Only the specific fragments used for annotation are shown.

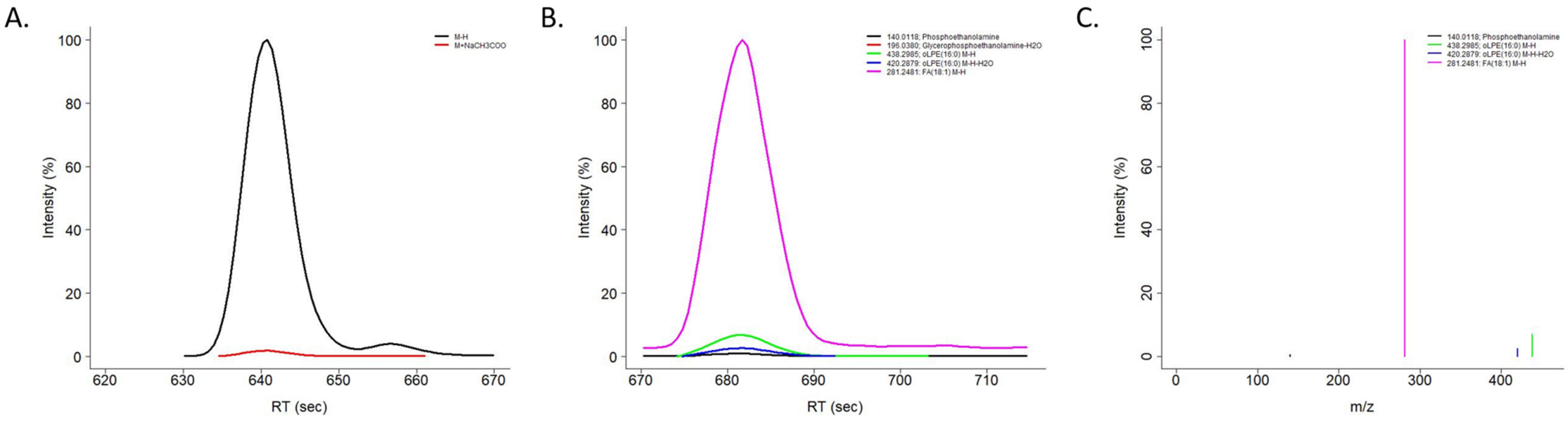

**Supplementary Figure S13. The PE(P-18:0/22:6) fragmentation profile for ESI+.** A) Chromatographic profiles of the precursor ions in MS1. B) Chromatographic profiles of the informative fragments in MS2 using DIA (35eV). Co-elution between the precursor and product fragments is observed (A-B). C) Fragmentation spectra for MS2 using DDA (35eV) for precursor ion 776.5594 (M+H). Only the specific fragments used for annotation are shown.

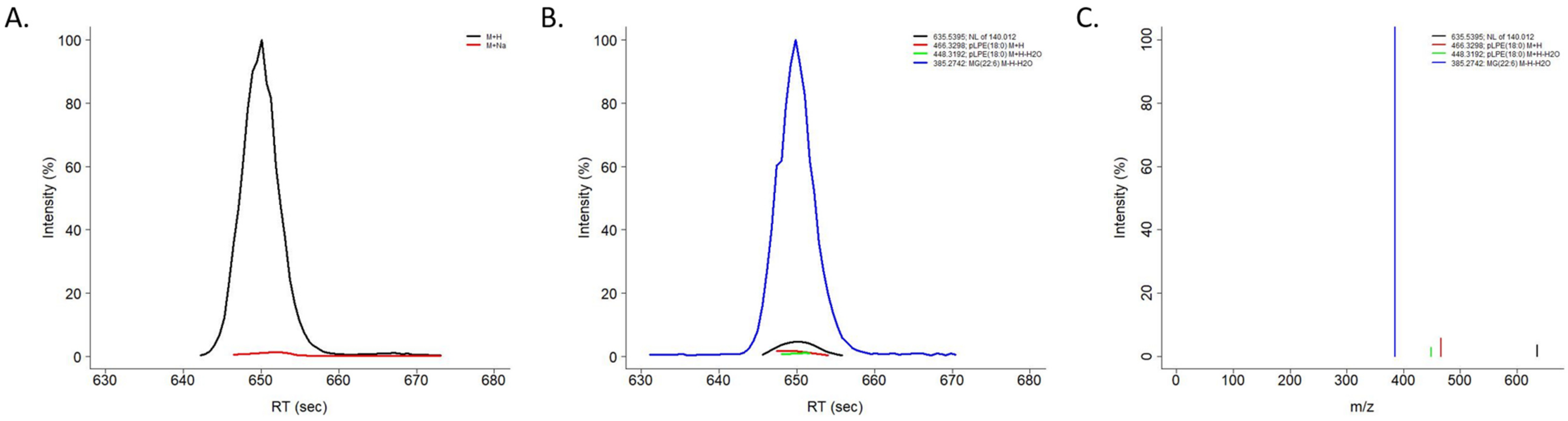

**Supplementary Figure S14. The PE(P-18:0/22:6) fragmentation profile for ESI-.** A) Chromatographic profiles of the precursor ions in MS1. B) Chromatographic profiles of the informative fragments in MS2 using DIA (35eV). Co-elution between the precursor and product fragments is observed (A-B). C) Fragmentation spectra for MS2 using DDA (35eV) for precursor ion 774.5438 (M-H). Only the specific fragments used for annotation are shown.

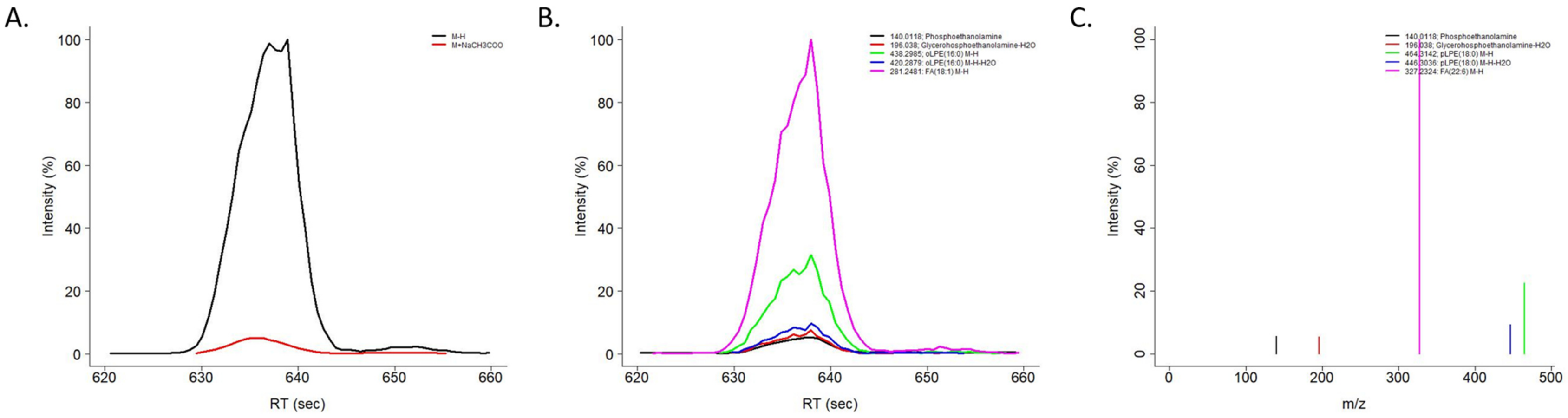

**Supplementary Figure S15. Web application screenshots.** The first five LipidMS web application tabs (A-E) take users through the complete data processing workflow. The last tab, "Resources" (F), contains links to access the extra documentation, examples and source code of the package.

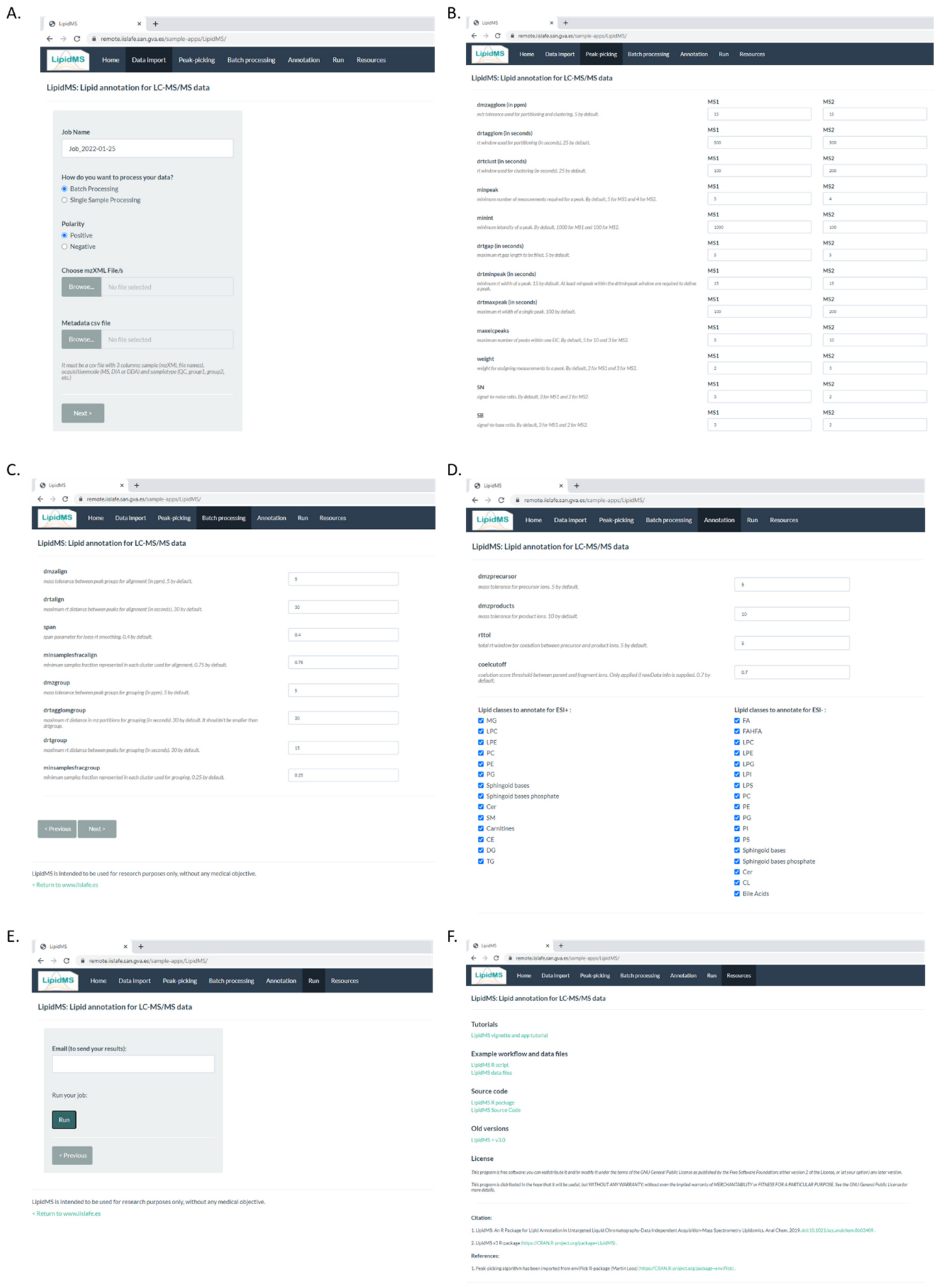

**Supplementary Figure S16. Summary of lipid annotations provided by LipidMS and MS-DIAL for the human serum pool.** Levels of structural information provided: Class, the detected fragments allow to provide information only about the total number of carbons and double bonds and the lipid class; FA, the identity of the fatty acyl moieties can be identified; FAposition, the actual position of the fatty acyl moieties within the lipid structure can be deduced.

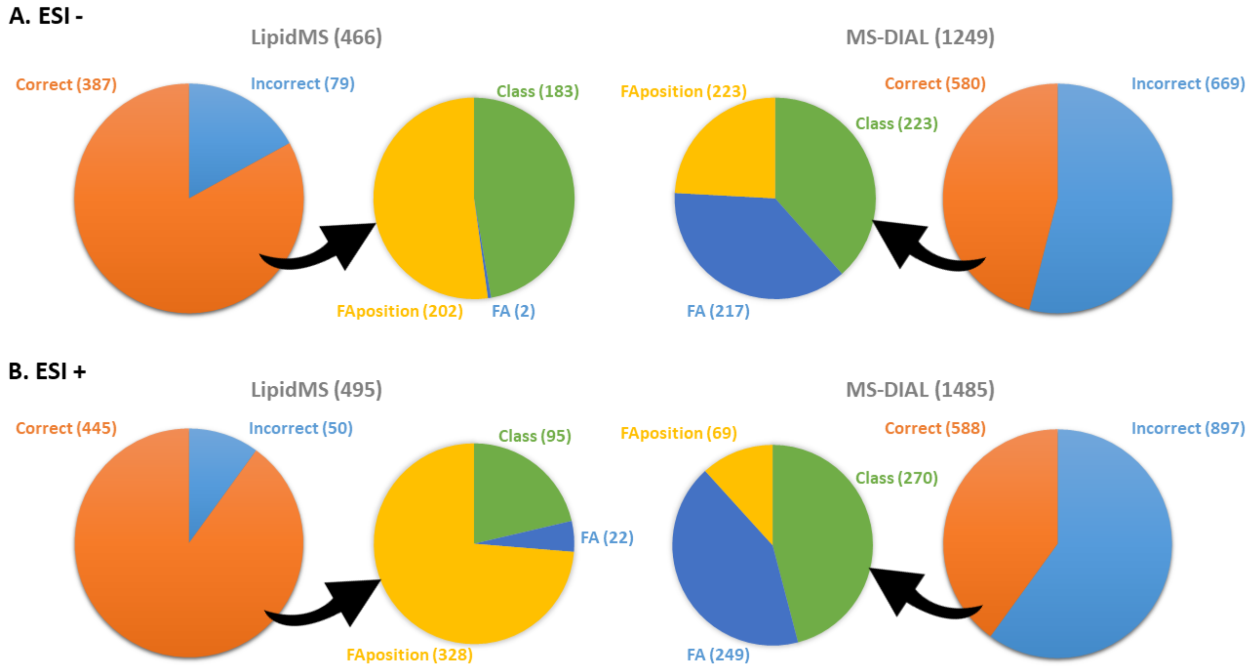

**Supplementary Tables**

**Supplementary Table S1. Adducts employed for the precursor identification of the new lipid classes.**

| Class [a] | ESI + | ESI - |
|---|---|---|
| **AcylCer** | M+H, M+H-H2O, M+Na | M-H, M+CH3COO |
| **CerP** | M+H | M-H |
| **oPC** | M+H, M+Na | M+CH3COO, M-CH3, M+CH3COO-CH3 |
| **pPC** | M+H, M+Na | M+CH3COO, M-CH3, M+CH3COO-CH3 |
| **oPE** | M+H, M+Na | M-H, M+NaCH3COO |
| **pPE** | M+H, M+Na | M-H, M+NaCH3COO |

AcylCer (acylceramides), CerP (ceramides phosphate), oPC (plasmanyl glycerophosphocholines), pPC (plasmenyl glycerophosphocholines), oPE (plasmanyl glycerophosphoethanolamines), pPE (plasmenyl glycerophosphoethanolamines).

**Supplementary Table S2. Fragmentation rules employed for the annotation of the new lipid classes in ESI+.**

| Class [a] | Class fragments (Subclass level) | Chain fragments (FA level) | Intensity rules (FA position level) |
|---|---|---|---|
| **AcylCer** | - | NL of acyl chain (Cer as M+H, M+H-$H_2O$ or M+H-2$H_2O$) Sph as M+H-$H_2O$ or M+H-2$H_2O$ FA as M+H | - |
| **CerP** | NL of phosphate group and $H_2O$ or 2 $H_2O$ (Cer as M+H-$H_2O$ or M+H-2$H_2O$) | Sph as M+H-2$H_2O$ Precursor - Sph | - |
| **oPC** | 104.1075, 184.0739, NL of 183.06604 | Sn1: oLPC as M+H or M+H-$H_2O$ Sn2: LPC as M+H-$H_2O$ or Precursor - sn1 | oLPC sn1 > 2 * LPC sn2 |
| **pPC** | 104.1075, 184.0739, NL of 183.06604 | Sn1: pLPC as M+H or M+H-$H_2O$ Sn2: LPC as M+H-$H_2O$ or Precursor - sn1 | pLPC sn1 > 2 * LPC sn2 |
| **oPE** | NL of 140.012 | Sn1: oLPE as M+H or M+H-$H_2O$ Sn2: MG as M+H-$H_2O$ | oLPE sn1 > 2 * MG sn2 |
| **pPE** | NL of 140.012 | Sn1: pLPE as M+H or M+H-$H_2O$ Sn2: MG as M+H-$H_2O$ | pLPE sn1 > 2 * MG sn2 |

AcylCer (acylceramides), NL (neutral loss), Sph (sphingoid base), FA (fatty acyl chain), CerP (ceramides phosphate), oPC (plasmanyl glycerophosphocholines), oLPC (plasmanyl lysoglycerophosphocholines), LPC (lysoglycerophosphocholines), pPC (plasmenyl glycerophosphocholines), pLPC (plasmenyl lysoglycerophosphocholines), oPE (plasmanyl glycerophosphoethanolamines), oLPE (plasmanyl lysoglycerophosphoethanolamines), LPE (lysoglycerophosphoethanolamines), MG (monoacylglycerol), pPE (plasmenyl glycerophosphoethanolamines), pLPE (plasmenyl lysoglycerophosphoethanolamines).

**Supplementary Table S3. Fragmentation rules employed for the annotation of the new lipid classes in ESI-.**

| Class [a] | Class fragments (Subclass level) | Chain fragments (FA level) | Intensity rules (FA position level) |
|---|---|---|---|
| **AcylCer** | - | NL of acyl chain (Cer as M-H)<br>NL of Sph (partial)<br>FA as M-H | Acyl chain > 5 * NL of Sph > 2 * FA |
| **CerP** | 78.9585, 96.9691 | SphP as M-H<br>FA as M-H | - |
| **oPC** | 168.0426, 224.0688, NL of $CH_3$ | Sn1: oLPC as M-$CH_3$ or M-$CH_3$-$H_2O$<br>Sn2: FA as M-H or M-$CO_2$-H | FA sn2 > 3 * oLPC sn1 |
| **pPC** | 168.0426, 224.0688, NL of $CH_3$ | Sn1: pLPC as M-$CH_3$ or M-$CH_3$-$H_2O$<br>Sn2: FA as M-H or M-$CO_2$-H | FA sn2 > 3 * pLPC sn1 |
| **oPE** | 140.0115, 196.038, 214.048 | Sn1: oLPE as M-H or M-H-$H_2O$<br>Sn2: FA as M-H | FA sn2 > 3 * oLPE sn1 |
| **pPE** | 140.0115, 196.038, 214.048 | Sn1: pLPE as M-H or M-H-$H_2O$<br>Sn2: FA as M-H | FA sn2 > 3 * pLPE sn1 |

AcylCer (acylceramides), NL (neutral loss), Sph (sphingoid base), FA (fatty acyl chain), CerP (ceramides phosphate), oPC (plasmanyl glycerophosphocholines), oLPC (plasmanyl lysoglycerophosphocholines), pPC (plasmenyl glycerophosphocholines), pLPC (plasmenyl lysoglycerophosphocholines), oPE (plasmanyl glycerophosphoethanolamines), oLPE (plasmanyl lysoglycerophosphoethanolamines), pPE (plasmenyl glycerophosphoethanolamines), pLPE (plasmenyl lysoglycerophosphoethanolamines).

**Supplementary Table S4. Summary of the expected lipid standard features detected and identified for each software package.**

| | Expected lipid standards | Expected lipid standard features | Total number of features | | | Expected features found | | | Significant features detected (adj. P.val < 0.05 and FOC > 2) | | | Identified features | | Identified lipid standards | |
|---|---|---|---|---|---|---|---|---|---|---|---|---|---|---|---|
| | | | **LipidMS** | **XCMS** | **MS-DIAL** | **LipidMS** | **XCMS** | **MS-DIAL** | **LipidMS** | **XCMS** | **MS-DIAL** | **LipidMS** | **MS-DIAL** | **LipidMS** | **MS-DIAL** |
| **ESI -** | 56 | 69 | 10220 | 19776 | 39366 | 68/69 | 56/69 | 56/69 | 59/69 | 56/69 | 54/69 | 55/69 | 43/69 | 48/56 | 42/56 |
| **ESI +** | 29 | 60 | 15354 | 24486 | 35897 | 54/60 | 58/60 | 59/60 | 41/60 | 46/60 | 42/60 | 43/60 | 33/60 | 22/29 | 21/29 |
| **Total** | 68 | 129 | 25574 | 44262 | 75263 | 122/129 | 114/129 | 115/129 | 100/129 | 102/129 | 96/129 | 98/129 | 76/129 | 60/68 | 56/68 |

### Supplementary Table S5. Lipid standards detected and identified in ESI-.

| Compound | Adduct | mz | RT (s) | Expected features found | | | Significant features detected (adj. P.val < 0.05 and FOC > 2) | | | Lipid standards identified | |
|---|---|---|---|---|---|---|---|---|---|---|---|
| | | | | LipidMS | XCMS | MS-DIAL | LipidMS | XCMS | MS-DIAL | LipidMS | MS-DIAL |
| **FA(10:0)** | M-H | 171.1385 | 54 | yes | yes | yes | yes | yes | yes | - | - |
| **FA(12:0)** | M-H | 199.1698 | 74 | yes | yes | yes | yes | yes | yes | FA(12:0) | FA(12:0) |
| **FA(14:0)** | M-H | 227.2011 | 116 | yes | yes | yes | yes | yes | yes | FA(14:0) | FA(14:0) |
| **FA(14:1)** | M-H | 225.1855 | 83 | yes | yes | yes | yes | yes | yes | FA(14:1) | - |
| **FA(15:0)** | M-H | 241.2167 | 149 | yes | yes | yes | yes | yes | yes | FA(15:0) | - |
| **FA(16:0)** | M-H | 255.2324 | 195 | yes | yes | yes | no | no | no | - | FA(16:0) |
| **FA(16:1)** | M-H | 253.2167 | 133 | yes | yes | yes | yes | yes | yes | FA(16:1) | FA(16:1) |
| **FA(17:0)** | M-H | 269.2481 | 245 | yes | yes | yes | yes | yes | yes | FA(17:0) | FA(17:0) |
| **FA(18:0)** | M-H | 283.2637 | 291 | yes | yes | yes | no | no | no | FA(18:0) | FA(18:0) |
| **FA(18:1)n9** | M-H | 281.2481 | 218 | yes | yes | yes | no | no | no | FA(18:1) | FA(18:1) |
| **FA(18:1)n7t** | M-H | 281.2481 | 229 | yes | yes | yes | yes | yes | yes | FA(18:1) | FA(18:1) |
| **FA(18:2)** | M-H | 279.2324 | 170 | yes | yes | yes | yes | yes | yes | FA(18:2) | FA(18:2) |
| **FA(18:3)n3** | M-H | 277.2168 | 111 | yes | no | yes | yes | - | no | - | - |
| **FA(18:3)n6** | M-H | 277.2168 | 115 | yes | yes | yes | yes | yes | yes | FA(18:3) | FA(18:3) |
| **FA(19:0)** | M-H | 297.2799 | 322 | yes | yes | yes | yes | yes | yes | - | FA(19:0) |
| **FA(20:0)** | M-H | 311.295 | 347 | yes | yes | yes | yes | yes | yes | FA(20:0) | FA(20:0) |
| **FA(20:1)** | M-H | 309.2794 | 302 | yes | yes | yes | yes | yes | yes | FA(20:1) | - |
| **FA(20:2)** | M-H | 307.2637 | 241 | yes | yes | yes | yes | yes | yes | FA(20:2) | FA(20:2) |
| **FA(20:3)** | M-H | 305.2481 | 182 | yes | yes | yes | yes | yes | yes | FA(20:3) | FA(20:3) |
| **FA(20:3)** | M-H | 305.2481 | 198 | yes | yes | yes | yes | yes | yes | FA(20:3) | FA(20:3) |
| **FA(20:4)n3** | M-H | 303.2324 | 132 | yes | no | yes | yes | - | yes | - | - |
| **FA(20:4)n6** | M-H | 303.2324 | 139 | yes | yes | yes | yes | yes | yes | FA(20:4) | FA(20:4) |
| **FA(20:5)** | M-H | 301.2168 | 103 | yes | yes | yes | yes | yes | yes | FA(20:5) | FA(20:5) |
| **FA(22:0)** | M-H | 339.3263 | 406 | yes | yes | yes | yes | yes | yes | FA(22:0) | FA(22:0) |
| **FA(22:1)** | M-H | 337.3107 | 352 | yes | yes | yes | yes | yes | yes | FA(22:1) | FA(22:1) |
| **FA(22:2)** | M-H | 335.295 | 315 | yes | yes | yes | yes | yes | yes | FA(22:2) | FA(22:2) |
| **FA(22:3)** | M-H | 333.2794 | 270 | yes | yes | yes | yes | yes | yes | FA(22:3) | FA(22:3) |
| **FA(22:4)n3** | M-H | 331.2637 | 212 | yes | yes | yes | yes | yes | no | - | - |
| **FA(22:4)n6** | M-H | 331.2637 | 217 | yes | no | yes | yes | - | yes | FA(22:4) | FA(22:4) |
| **FA(22:5)n3** | M-H | 329.2481 | 158 | yes | yes | yes | yes | yes | yes | FA(22:5) | FA(22:5) |
| **FA(22:5)n6** | M-H | 329.2481 | 165 | yes | yes | yes | yes | yes | yes | FA(22:5) | FA(22:5) |
| **FA(22:6)** | M-H | 327.2324 | 121 | yes | yes | yes | yes | yes | yes | FA(22:6) | FA(22:6) |
| **FA(24:0)** | M-H | 367.3576 | 483 | yes | yes | yes | yes | yes | yes | FA(24:0) | FA(24:0) |
| **FA(24:1)** | M-H | 365.3419 | 408 | yes | yes | yes | yes | yes | yes | FA(24:1) | FA(24:1) |
| **FA(26:0)** | M-H | 395.3889 | 585 | yes | yes | yes | yes | yes | yes | FA(26:0) | FA(26:0) |
| **LPC(17:0)** | M+CH3COO | 568.3621 | 135 | yes | yes | yes | yes | yes | yes | LPC(17:0) | LPC(17:0) |
| **LPC(17:0)** | M-CH3 | 494.3247 | 135 | yes | yes | no | yes | yes | - | LPC(17:0) | - |
| **PC(17:0/17:0)** | M+CH3COO | 820.6074 | 637 | yes | yes | no | yes | yes | - | PC(17:0/17:0) | - |
| **PC(17:0/17:0)** | M-CH3 | 746.57 | 637 | yes | yes | yes | yes | yes | yes | PC(17:0/17:0) | *PE(18:0_18:0) |
| **PC(16:0/18:1)** | M+CH3COO | 818.5917 | 544 | yes | yes | yes | no | no | no | PC(16:0/18:1) | *PC(O-17:0_17:2) |
| **PC(16:0/18:1)** | M-CH3 | 744.5544 | 544 | yes | yes | yes | no | no | no | PC(16:0/18:1) | - |
| **PC(18:0/18:2)** | M+CH3COO | 844.6074 | 567 | yes | yes | yes | no | no | no | PC(18:0/18:2) | - |
| **PC(18:0/18:2)** | M-CH3 | 770.57 | 567 | yes | yes | yes | no | no | no | PC(18:0/18:2) | *CL(18:0_18:0_22:0_20:3) |
| **PC(O-16:0/20:5)** | M+CH3COO | 824.5812 | 462 | yes | yes | yes | yes | yes | yes | PC(O-16:0/20:5) | PC(O-16:0_20:5) |
| **PC(O-16:0/20:5)** | M-CH3 | 750.5438 | 462 | yes | yes | yes | yes | yes | yes | - | *PE(O-18:0_20:5) |
| **PC(P-18:0/20:4)** | M+CH3COO | 852.6125 | 595 | yes | yes | yes | yes | yes | yes | PC(P-18:0/20:4) | *PC(O-18:1_20:4) |
| **PC(P-18:0/20:4)** | M-CH3 | 778.5751 | 595 | no | no | no | - | - | - | - | - |
| **PE(17:0/17:0)** | M-H | 718.5387 | 659 | yes | yes | yes | yes | yes | yes | - | PE(17:0_17:0) |
| **PE(16:0/18:1)** | M-H | 716.5231 | 561 | yes | yes | yes | yes | yes | yes | PE(16:0/18:1) | PE(16:0_18:1) |
| **PE(O-16:0/18:1)** | M-H | 702.5438 | 630 | yes | yes | yes | yes | yes | yes | - | PE(O-16:0_18:1) |
| **PE(P-18:0/22:6)** | M-H | 774.5438 | 588 | yes | yes | yes | yes | yes | yes | PE(P-18:0/22:6) | *PE(O-18:1_22:6) |
| **PG(17:0/17:0)** | M-H | 749.5333 | 480 | yes | yes | yes | yes | yes | yes | PG(17:0/17:0) | PG(17:0_17:0) |
| **PG(16:0/18:1)** | M-H | 747.5177 | 426 | yes | yes | yes | yes | yes | yes | PG(16:0/18:1) | PG(16:0_18:1) |
| **PI(17:0/14:1)** | M-H | 793.4867 | 350 | yes | yes | yes | yes | yes | yes | PI(17:0/14:1) | PI(17:0_14:1) |
| **PS(17:0/17:0)** | M-H | 762.5285 | 473 | yes | yes | yes | yes | yes | yes | PS(17:0/17:0) | PS(17:0_17:0) |
| **PS(17:0/17:0)** | M+Na-2H | 784.5099 | 473 | yes | yes | yes | yes | yes | yes | - | - |
| **PS(16:0/18:1)** | M-H | 760.5129 | 420 | yes | yes | yes | yes | yes | yes | PS(16:0_18:1) | PS(16:0_18:1) |
| **PS(16:0/18:1)** | M+Na-2H | 782.4943 | 420 | yes | yes | no | yes | yes | - | - | - |
| **CL(18:1/18:1/18:1/18:1)** | M-H | 1456.027 | 1014 | yes | yes | yes | yes | yes | yes | CL(18:1/18:1/18:1/18:1) | CL(36:2_36:2) |
| **CL(18:1/18:1/18:1/18:1)** | M+Na-2H | 1478.009 | 1014 | yes | yes | yes | yes | yes | yes | CL(18:1/18:1/18:1/18:1) | *CL(37:2_37:5) |
| **SM(d18:1/17:0)** | M+CH3COO | 775.5972 | 476 | yes | yes | yes | yes | yes | yes | SM(18:1/17:0) | SM(d18:1/17:0) |
| **SM(d18:1/17:0)** | M-CH3 | 701.5598 | 476 | yes | yes | yes | yes | yes | yes | SM(35:1) | - |
| **SM(d18:1/16:0)** | M+CH3COO | 761.5815 | 440 | yes | yes | yes | no | no | no | SM(18:1/16:0) | SM(d18:1/16:0) |
| **SM(d18:1/16:0)** | M-CH3 | 687.5441 | 440 | yes | yes | yes | no | no | no | SM(18:1/16:0) | - |
| **Cer(d18:1/17:0)** | M+CH3COO | 550.5199 | 603 | yes | yes | yes | yes | yes | yes | Cer(d18:1/17:0) | Cer(d18:1/17:0) |
| **Cer(d18:1/17:0)** | M-CH3 | 610.5416 | 603 | yes | yes | yes | yes | yes | yes | Cer(d18:1/17:0) | Cer(d18:1/17:0) |
| **CerP(d18:1/16:0)** | M-H | 616.4707 | 384 | yes | yes | yes | yes | yes | yes | CerP(d34:1) | *SL(15:1;O/19:0;O) |
| **AcylCer(18:1-d18:1/17:0)** | M-H | 814.7652 | 1030 | yes | yes | yes | yes | yes | yes | - | - |
| **AcylCer(18:1-d18:1/17:0)** | M+CH3COO | 874.7869 | 1030 | yes | yes | yes | yes | yes | yes | - | - |

### Supplementary Table S6. Lipid standards detected and identified in ESI+.

| Compound | Adduct | mz | RT (s) | Expected features found | | | Significant features detected (adj. P.val < 0.05 and FOC > 2) | | | Lipid standards identified | |
|---|---|---|---|---|---|---|---|---|---|---|---|
| | | | | LipidMS | XCMS | MS-DIAL | LipidMS | XCMS | MS-DIAL | LipidMS | MS-DIAL |
| **LPC(17:0)** | M+H | 510.356 | 142 | yes | yes | yes | yes | yes | yes | LPC(17:0) | LPC(17:0) |
| **LPC(17:0)** | M+Na | 532.3372 | 142 | yes | yes | yes | yes | yes | yes | LPC(17:0) | *LPC(19:3) |
| **PC(17:0/17:0)** | M+H | 762.6013 | 730 | yes | yes | yes | yes | yes | no | PC(17:0/17:0) | PC(17:0_17:0) |
| **PC(17:0/17:0)** | M+Na | 784.5827 | 730 | yes | yes | yes | yes | yes | yes | - | - |
| **PC(16:0/18:1)** | M+H | 760.5856 | 609 | yes | yes | yes | no | no | no | PC(16:0/18:1) | PC(16:0_18:1) |
| **PC(16:0/18:1)** | M+Na | 782.567 | 609 | yes | yes | yes | no | no | no | PC(16:0/18:1) | PC(34:1) |
| **PC(18:0/18:2)** | M+H | 786.6013 | 640 | yes | yes | yes | no | no | no | PC(18:0/18:2) | PC(18:0_18:2) |
| **PC(18:0/18:2)** | M+Na | 808.5827 | 640 | yes | yes | yes | no | no | no | *PC(18:1/18:1) | PC(36:2) |
| **PC(O-16:0/20:5)** | M+H | 766.5751 | 505 | yes | yes | yes | yes | yes | yes | PC(O-16:0/20:5) | PC(O-36:5) |
| **PC(O-16:0/20:5)** | M+Na | 788.5565 | 505 | yes | yes | yes | yes | yes | yes | PC(O-16:0/20:5) | - |
| **PC(P-18:0/20:4)** | M+H | 794.6064 | 675 | yes | yes | yes | yes | yes | yes | PC(P-18:0/20:4) | *PC(O-38:5) |
| **PC(P-18:0/20:4)** | M+Na | 816.5878 | 675 | yes | yes | yes | yes | yes | no | PC(P-18:0/20:4) | *PC(O-40:8) |
| **PE(17:0/17:0)** | M+H | 720.5543 | 762 | yes | yes | yes | yes | yes | yes | PE(17:0/17:0) | PE(17:0/17:0) |
| **PE(17:0/17:0)** | M+Na | 742.5357 | 762 | no | no | no | - | - | - | - | - |
| **PE(16:0/18:1)** | M+H | 718.5387 | 635 | yes | yes | yes | yes | yes | yes | - | - |
| **PE(16:0/18:1)** | M+Na | 740.5201 | 635 | no | yes | yes | - | yes | yes | - | - |
| **PE(O-16:0/18:1)** | M+H | 704.5594 | 725 | yes | yes | yes | yes | yes | yes | PE(O-16:0/18:1) | PE(O-34:1) |
| **PE(O-16:0/18:1)** | M+Na | 726.5408 | 725 | no | yes | yes | - | yes | yes | - | - |
| **PE(P-18:0/22:6)** | M+H | 776.5594 | 675 | yes | yes | yes | yes | yes | yes | *PE(O-18:1_22:6) | PE(P-18:0_22:6) |
| **PE(P-18:0/22:6)** | M+Na | 798.5408 | 675 | no | yes | yes | - | yes | no | - | - |
| **PG(17:0/17:0)** | M+H | 751.5489 | 587 | yes | yes | yes | no | yes | yes | - | - |
| **PG(17:0/17:0)** | M+Na | 773.5303 | 587 | no | no | yes | - | - | yes | - | - |
| **PG(16:0/18:1)** | M+H | 749.5333 | 503 | yes | yes | yes | yes | yes | yes | - | - |
| **PG(16:0/18:1)** | M+Na | 771.5147 | 503 | yes | yes | yes | yes | yes | yes | - | - |
| **PI(17:0/14:1)** | M+H | 795.5023 | 384 | yes | yes | yes | yes | yes | yes | - | - |
| **PI(17:0/14:1)** | M+NH4 | 812.5289 | 384 | yes | yes | yes | yes | yes | yes | PI(31:1) | PI(31:1) |
| **PI(17:0/14:1)** | M+Na | 817.4837 | 384 | yes | yes | yes | yes | yes | yes | PI(31:1) | - |
| **SM(d18:1/17:0)** | M+H | 717.5911 | 520 | yes | yes | yes | yes | yes | yes | SM(d18:1/17:0) | *SM(d17:1/18:0) |
| **SM(d18:1/17:0)** | M+Na | 739.5725 | 520 | yes | yes | yes | yes | yes | yes | SM(d35:1) | SM(d35:1) |
| **SM(d18:1/16:0)** | M+H | 703.5754 | 476 | yes | yes | yes | no | no | no | SM(d18:1/16:0) | SM(d18:1/16:0) |
| **SM(d18:1/16:0)** | M+Na | 725.5568 | 476 | yes | yes | yes | no | no | no | SM(d18:1/16:0) | SM(d34:1) |
| **Cer(d18:1/17:0)** | M+H | 552.5355 | 682 | yes | yes | yes | yes | yes | yes | Cer(d18:1/17:0) | Cer(d18:1/17:0) |
| **Cer(d18:1/17:0)** | M+H-H2O | 534.5249 | 682 | yes | yes | yes | yes | yes | yes | Cer(d18:1/17:0) | Cer(d18:1/17:0) |
| **Cer(d18:1/17:0)** | M+Na | 574.5169 | 682 | yes | yes | yes | yes | yes | no | Cer(d18:1/17:0) | *Cer(d18:1/19:2) |
| **CerP(d18:1/16:0)** | M+H | 618.4863 | 377 | yes | yes | yes | no | no | no | - | - |
| **AcylCer(18:1;18:1/17:0)** | M+H | 816.7808 | 1077 | yes | yes | yes | yes | yes | yes | AcylCer(18:1;d18:1/17:0) | *Cer(53:3;3O) |
| **AcylCer(18:1;18:1/17:0)** | M+H-H2O | 798.7702 | 1077 | yes | yes | yes | yes | yes | yes | AcylCer(18:1;d18:1/17:0) | *Cer(53:3;3O) |
| **AcylCer(18:1;18:1/17:0)** | M+Na | 838.7622 | 1077 | yes | yes | yes | yes | yes | yes | AcylCer(18:1;d18:1/17:0) | *Cer(55:6;3O) |
| **MG(17:0)** | M+Na | 367.2818 | 278 | yes | yes | yes | yes | yes | yes | - | - |
| **DG(17:0/17:0)** | M+H-H2O | 579.5351 | 953 | yes | yes | yes | yes | yes | yes | DG(17:0/17:0) | - |
| **DG(17:0/17:0)** | M+NH4 | 614.5723 | 953 | yes | yes | yes | yes | yes | yes | DG(17:0/17:0) | DG(17:0_17:0) |
| **DG(17:0/17:0)** | M+Na | 619.5271 | 953 | yes | yes | yes | yes | yes | yes | DG(17:0/17:0) | DG(34:0) |
| **TG(8:0/8:0/8:0)** | M+NH4 | 488.3951 | 391 | yes | yes | yes | yes | yes | yes | TG(8:0/8:0/8:0) | TG(8:0_8:0_8:0) |
| **TG(8:0/8:0/8:0)** | M+Na | 493.3499 | 391 | yes | yes | yes | yes | yes | yes | TG(8:0/8:0/8:0) | TG(8:0_8:0_8:0) |
| **TG(10:0/10:0/10:0)** | M+NH4 | 572.489 | 648 | yes | yes | yes | yes | yes | yes | TG(10:0/10:0/10:0) | TG(10:0_10:0_10:0) |
| **TG(10:0/10:0/10:0)** | M+Na | 577.4438 | 648 | yes | yes | yes | yes | yes | yes | TG(10:0/10:0/10:0) | TG(10:0_10:0_10:0) |
| **TG(12:0/12:0/12:0)** | M+NH4 | 656.5829 | 966 | yes | yes | yes | yes | yes | yes | TG(12:0/12:0/12:0) | TG(12:0_12:0_12:0) |
| **TG(12:0/12:0/12:0)** | M+Na | 661.5376 | 966 | yes | yes | yes | yes | yes | yes | TG(12:0/12:0/12:0) | TG(12:0_12:0_12:0) |
| **TG(14:0/14:0/14:0)** | M+NH4 | 740.6767 | 1063 | yes | yes | yes | yes | yes | yes | TG(14:0/14:0/14:0) | TG(14:0_14:0_14:0) |
| **TG(14:0/14:0/14:0)** | M+Na | 745.6315 | 1063 | yes | yes | yes | yes | yes | yes | TG(14:0/14:0/14:0) | TG(14:0_14:0_14:0) |
| **TG(16:0/16:0/16:0)** | M+NH4 | 824.7707 | 1140 | yes | yes | yes | no | no | no | TG(16:0/16:0/16:0) | TG(16:0_16:0_16:0) |
| **TG(16:0/16:0/16:0)** | M+Na | 829.7255 | 1140 | yes | yes | yes | yes | yes | yes | TG(16:0/16:0/16:0) | - |
| **TG(17:0/17:0/17:0)** | M+NH4 | 866.8176 | 1173 | yes | yes | yes | yes | yes | yes | TG(17:0/17:0/17:0) | TG(17:0_17:0_17:0) |
| **TG(17:0/17:0/17:0)** | M+Na | 871.7724 | 1173 | yes | yes | yes | yes | yes | yes | TG(17:0/17:0/17:0) | TG(17:0_17:0_17:0) |
| **TG(18:1/16:0/18:1)** | M+NH4 | 876.802 | 1142 | yes | yes | yes | no | no | no | TG(18:1/16:0/18:1) | TG(16:0_18:1_18:1) |
| **TG(18:1/16:0/18:1)** | M+Na | 881.7568 | 1142 | yes | yes | yes | no | no | no | TG(18:1/16:0/18:1) | TG(16:0_18:1_18:1) |
| **TG(16:0/16:0/18:1)** | M+NH4 | 850.7863 | 1140 | yes | yes | yes | no | no | no | TG(16:0/16:0/18:1) | TG(16:0_16:0_18:1) |
| **TG(16:0/16:0/18:1)** | M+Na | 855.7411 | 1140 | yes | yes | yes | no | no | no | TG(16:0/16:0/18:1) | TG(16:0_16:0_18:1) |
| **CE(17:0)** | 2M+NH4 | 1295.235 | 1157 | yes | yes | yes | yes | yes | no | - | - |
| **CE(17:0)** | 2M+Na | 1300.189 | 1157 | no | yes | yes | - | yes | yes | - | - |

**Supplementary Table S7. Summary of the identified lipids in ESI-.**

| | Total | Correct | Class | FA | FA position | Incorrect | Unique | Missing |
|---|---|---|---|---|---|---|---|---|
| **LipidMS** | 466 | 387 (83%) | 183 | 2 | 202 | 79 (17%) | 152 | 342 |
| **MSDial** | 1249 | 580 (46%) | 223 | 217 | 140 | 669 (54%) | 345 | 92 |

Total: total number of annotated lipids. Correct: lipids whose proposed annotation is correct based on the observed spectra. Class: specific subclass fragments are found, but only the total number of carbons and double bonds of the chains can be proposed based on the precursor ion. FA: the specific chain fragments that inform about the composition of fatty acyl chains are found. FA position: when the specific fatty acyl chain position can be elucidated based on chain fragments intensity ratios. Incorrect: lipids for which an incorrect annotation is provided, or non lipid features are annotated as lipids. Unique: lipids that are annotated exclusively by one of the software packages. Missing: the lipids with confirmed lipid identity, but are not annotated by one of the software packages.

**Supplementary Table S8. Summary of the identified lipids in ESI+.**

| | Total | Correct | Class | FA | FA position | Incorrect | Unique | Missing |
|---|---|---|---|---|---|---|---|---|
| **LipidMS** | 495 | 445 (90%) | 95 | 22 | 328 | 50 (10%) | 140 | 297 |
| **MSDial** | 1485 | 588 (40%) | 270 | 249 | 69 | 897 (60%) | 283 | 124 |

Total: total number of annotated lipids. Correct: lipids whose proposed annotation is correct based on the observed spectra. Class: specific subclass fragments are found, but only the total number of carbons and double bonds of the chains can be proposed based on the precursor ion. FA: the specific chain fragments that inform about the composition of fatty acyl chains are found. FA position: when the specific fatty acyl chain position can be elucidated based on chain fragments intensity ratios. Incorrect: lipids for which an incorrect annotation is provided, or non lipid features are annotated as lipids. Unique: lipids that are annotated exclusively by one of the software packages. Missing: the lipids with confirmed lipid identity, but are not annotated by one of the software packages.
